# WNT Signalling Promotes NF-κB Activation and Drug Resistance in KRAS-Mutant Colorectal Cancer

**DOI:** 10.1101/2023.12.21.572810

**Authors:** Bojie Cong, Evangelia Stamou, Kathryn Pennel, Molly Mckenzie, Amna Matly, Sindhura Gopinath, Joanne Edwards, Ross Cagan

**Author notes:** To whom correspondence should be addressed. Ross Cagan.

## Abstract

Approximately 40% of colorectal cancer (CRC) cases are characterized by KRAS mutations, rendering them insensitive to most CRC therapies. While the reasons for this resistance remain incompletely understood, one key aspect is genetic complexity: in CRC, oncogenic KRAS is most commonly paired with mutations that alter WNT and P53 activities (“RAP”). Here, we demonstrate that elevated WNT activity upregulates canonical (NF-κB) signalling in both *Drosophila* and human RAS mutant tumours. This upregulation required Toll-1 and Toll-9 and resulted in reduced efficacy of RAS pathway targeted drugs such as the MEK inhibitor trametinib. Inhibiting WNT activity pharmacologically significantly suppressed trametinib resistance in RAP tumours and more genetically complex RAP-containing ‘patient avatar’ models. WNT/MEK drug inhibitor combinations were further improved by targeting *brm*, *shg*, *ago*, *rhoGAPp190* and *upf1*, highlighting these genes as candidate biomarkers for patients sensitive to this duel approach. These findings shed light on how genetic complexity impacts drug resistance and proposes a therapeutic strategy to reverse this resistance.

## Introduction

Colorectal cancer (CRC) emerged as the second leading cause of cancer-related deaths worldwide. This disease is characterized by uncontrolled cell growth, primarily driven by a complex array of genetic mutations (*1–3*). Notably, CRC cases featuring RAS mutations have displayed a notable insensitivity to most targeted therapies for CRC in clinics (*4*). Despite the development of successive generations of inhibitors targeting the RAS pathway—which have demonstrated promise in pre-clinical studies—these inhibitors have shown minimal or transitory activity in RAS-mutant CRC patients. A better understanding of the characteristics of tumours in clinical contexts is needed to develop more durable treatments.

Genetic complexity represents a consistent and key clinical hallmark of tumours. Experiments in both Drosophila and mouse models have found that oncogenic KRAS mutations alone initiate benign tumours (*5*, *6*). The progression to malignancy requires acquisition of additional mutations in genes such as *P53* and the WNT regulator *APC* (*7*, *8*), which are frequently mutated together in human CRC (*9*, *10*). Our recent studies involving *Drosophila* CRC models have also demonstrated that genetic complexity amplifies metastatic potential and, key to this report, fosters drug resistance (*9*). The precise mechanisms by which genetic complexity influences drug response and patient health remain poorly understood.

Here, we examine the impact of mutated *apc* and *p53* on Ras^G12V^ tumours in *Drosophila*. We found that elevated WNT (Wg in *Drosophila*) activity led to upregulation of canonical nuclear factor-kappa B (NF-κB) signalling in Ras^G12V^ tumours, in turn promoting overgrowth of Ras^G12V^ tumours and reducing host survival. We also found that this elevation of canonical NF-κB led to emergent resistance to drugs such as the MEK inhibitor trametinib; this resistance was reversed by inhibiting WNT activity with compounds such as PNU-74654 or LF3, leading to strong trametinib-mediated rescue. Patient CRC samples with high WNT activity and oncogenic KRAS were associated with upregulation of canonical NF-κB signalling, consistent with our *Drosophila* findings and suggestive of a novel approach to the treatment of KRAS-mutant CRC.

## Results

### WNT signalling reduces trametinib efficacy by elevating canonical NF-κB signalling

Ras proteins regulate key cellular processes through multiple pathways including the canonical Raf-MEK-ERK (MAPK) signalling pathway. The FDA-approved drug trametinib is a potent and precise MEK inhibitor that, despite strong preclinical activity, has consistently failed to demonstrate significant efficacy in CRC patients (*11*, *12*). Consistent with these reports, we observed that trametinib—delivered orally in the food—strongly rescued tumour-induced lethality in *Drosophila* CRC models that target RAS^G12V^ to the hindgut (*byn>Ras^G12V^*; Figure 1A) but failed to rescue a multigenic *Ras^G12V^*, *Apc^RNAi^*, *P53^RNAi^* CRC model (*byn>RAP;* Figure 1A). Trametinib did not affect control animals (Supplemental Figure 1a). These data indicate an emergent resistance to trametinib in *byn>RAP* tumours.

**Figure 1:**
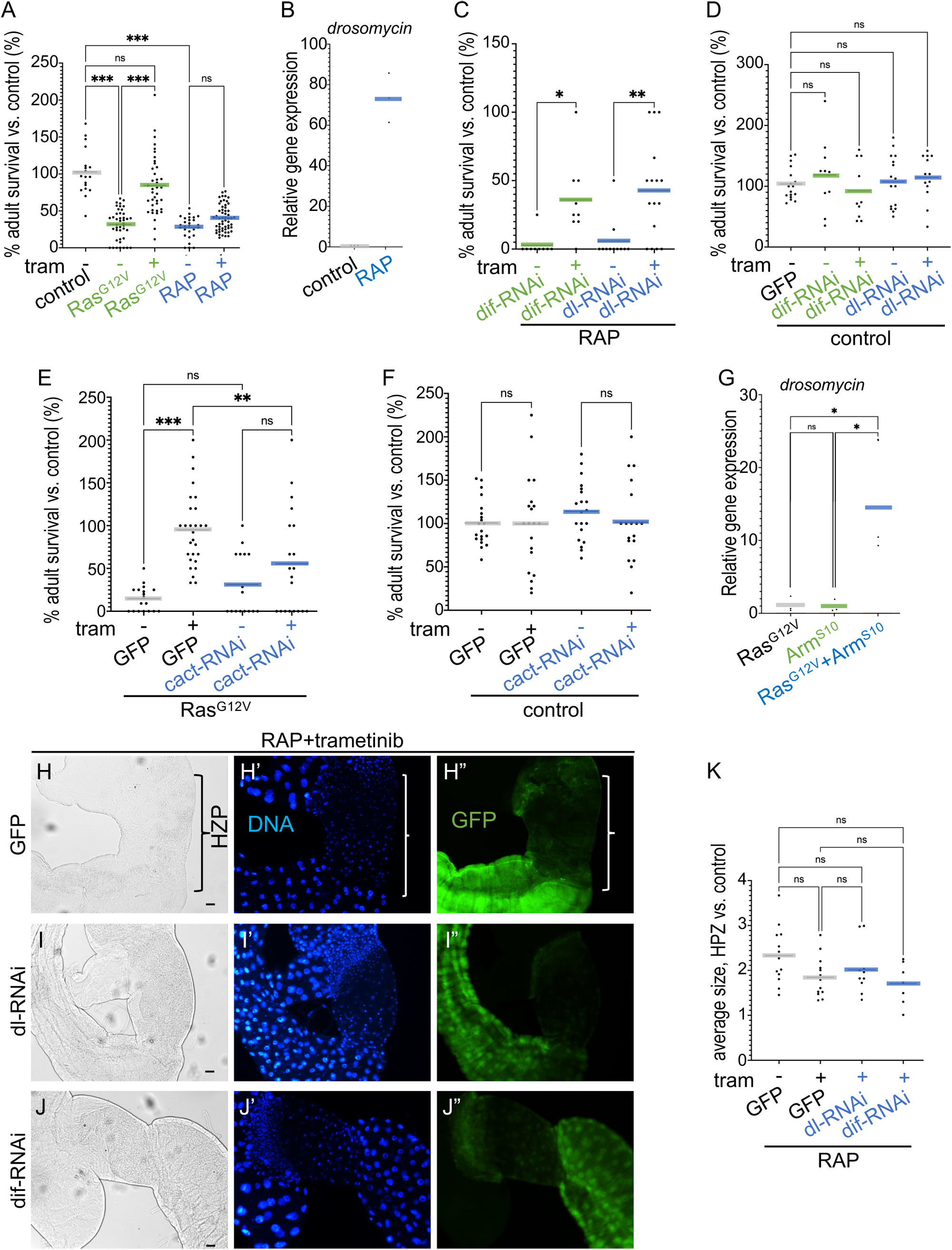
WNT signalling induced trametinib resistance through enhancing canonical NF-κB signalling. **(A, C, D, E, F)** Percent survival of transgenic flies to adulthood relative to control flies was quantified in the presence or absence of trametinib (1 μM). **(A)** Control, *Ras^G12V^*, and *RAP*; **(C)** *RAP* +*dif-RNAi* and *RAP* +*dl-RNAi*; **(D)** *dif-RNAi* and *dl-RNAi*; **(E)** *Ras^G12V^*+GFP (control) and *Ras^G12V^*+*cact-RNAi*; **(F)** GFP and *cact-RNAi*. **(B, G)** Expression levels of drosomycin were quantified for each genotype by quantitative RT-PCR. **(B)** Control and RAP; **(G)** *Ras^G12V^*, *Arm^S10^* and *Ras^G12V^*+*Arm^S10^*. **(H-J)** Images of the digestive tract of third instar larvae in the present of trametinib (1 μM), which include the hindgut proliferation zone (HPZ). Nuclei are visualized with 4′,6-diamidino-2-phenylindole (DAPI) staining, hindgut is visualized by GFP. Scale bar 100μm. **(K)** The average of hindgut proliferation zone (HPZ) size was measured by Fiji ImageJ and quantified as relative size to wild-type hindgut.

In a screen for pathways that distinguish RAS^G12V^ and RAP models (not shown), we identified differences in NF-κB pathway activity. Previous studies have shown that inhibition of NF-κB signalling increases sensitivity of HTC15 human colon cancer cells to the chemotherapeutic daunomycinby modulating drug uptake (*13*). Interestingly, we found that canonical NF-κB signalling—also known as the Toll pathway in *Drosophila—*was upregulated in *byn*>*RAP* tumours as visualized by the target gene *drosomycin* (Figure 1B) (*14*). Inhibition of canonical NF-κB signalling by targeted knockdown of NF-κB factors such as *dorsal-related immunity factor* (*dif*) or *dorsal* (*dl*) (*15*) significantly rescued tumour-induced lethality in the presence of trametinib (Figure 1C). Knockdown did not affect survival of control animals or *byn>RAP* animals in the absence of trametinib (Figure 1C, 1D). Conversely, elevating canonical NF-κB activity by targeted knockdown of the NF-κB inhibitor *cactus* (*cact*, an IκB orthologue) (*16*, *17*) reduced trametinib efficacy in *byn>Ras^G12V^*tumours; this elevation of canonical NF-κB activity did not affect lethality in the absence of trametinib or control animals (Figure 1E, 1F).

Our data suggested that cooperative activation of Wg and Ras led to an increase in canonical NF-κB pathway activity. Consistent with this view, the canonical NF-κB pathway reporter *drosomycin* was elevated when Ras and Wg activities were elevated together in the hindgut but not when either gene was activated alone (Figure 1G). These data indicate that WNT activity induces trametinib resistance at least in part by elevating canonical NF-κB signalling in *byn>Ras^G12V^* tumours.

To gain a deeper understanding of drug impact on *Ras^G12V^*vs. *RAP* tumours, we assessed the effect of trametinib on tumour growth in the *Drosophila* hindgut proliferative zone (HPZ). Trametinib exhibited a near complete suppression of HPZ tumour overgrowth in *byn>Ras^G12V^*tumours (Supplemental Figure 1b, 1e-1g). However, trametinib only partially suppressed HPZ tumour overgrowth of *byn>RAP* tumours (Figure 1H, 1K, Supplemental Figure 1k). Inhibition of canonical NF-κB activity by knockdown of *dl* or *dif* did not significantly suppress tumour overgrowth in the presence of trametinib in *byn>RAP* tumours (Figure 1I and 1J compared to 1H, quantified in 1K). Moreover, elevating Wg signalling did not enhance tumour overgrowth in *byn>Ras^G12V^* tumours or control animals (Supplemental Figure 1h and 1i compared to S1f and S1e, quantified in S1c). However, in Ras^G12V^ tumour, the overgrowth of HPZ was significantly enhanced by both Wg signalling activity and p53 defect, while p53 alone had a weaker effect on promoting overgrowth of HPZ (Supplemental Figure 1j and S1k compared to S1f, quantified in S1d). These data suggest that Wg activation and p53 defect promotes overgrowth in Ras^G12V^ tumour and canonical NF-κB signalling reduces trametinib efficacy by primarily modulating host survival rather than tumour growth in *Drosophila*.

### Toll-1 and Toll-9 are required for upregulation of NF-κB activity in RAP tumours

Our next question was to understand the mechanism by which Wg signalling upregulates canonical NF-κB activity in *byn>RAP* tumours. A recent study had unveiled that Toll-9 alone could effectively enhance canonical NF-κB activity in *Drosophila* imaginal discs, and this upregulation of NF-κB activity was reliant on Toll-1 (*18*). We observed that nuclear translocation of Dorsal was markedly increased in *byn>RAP* tumours compared to control (Figure 2B compared to Figure 2A). However, this upregulation of canonical NF-κB activity was suppressed by the knockdown of Toll-1 or Toll-9 in *byn>RAP* tumours (Figure 2C and 2D compared to Figure 2B). Furthermore, inhibition of canonical NF-κB activity by the knockdown of Toll-1 or Toll-9 significantly rescued tumour-induced lethality in the presence of trametinib, while it had no effect in the absent of trametinib (Figure 2E).

**Figure 2:**
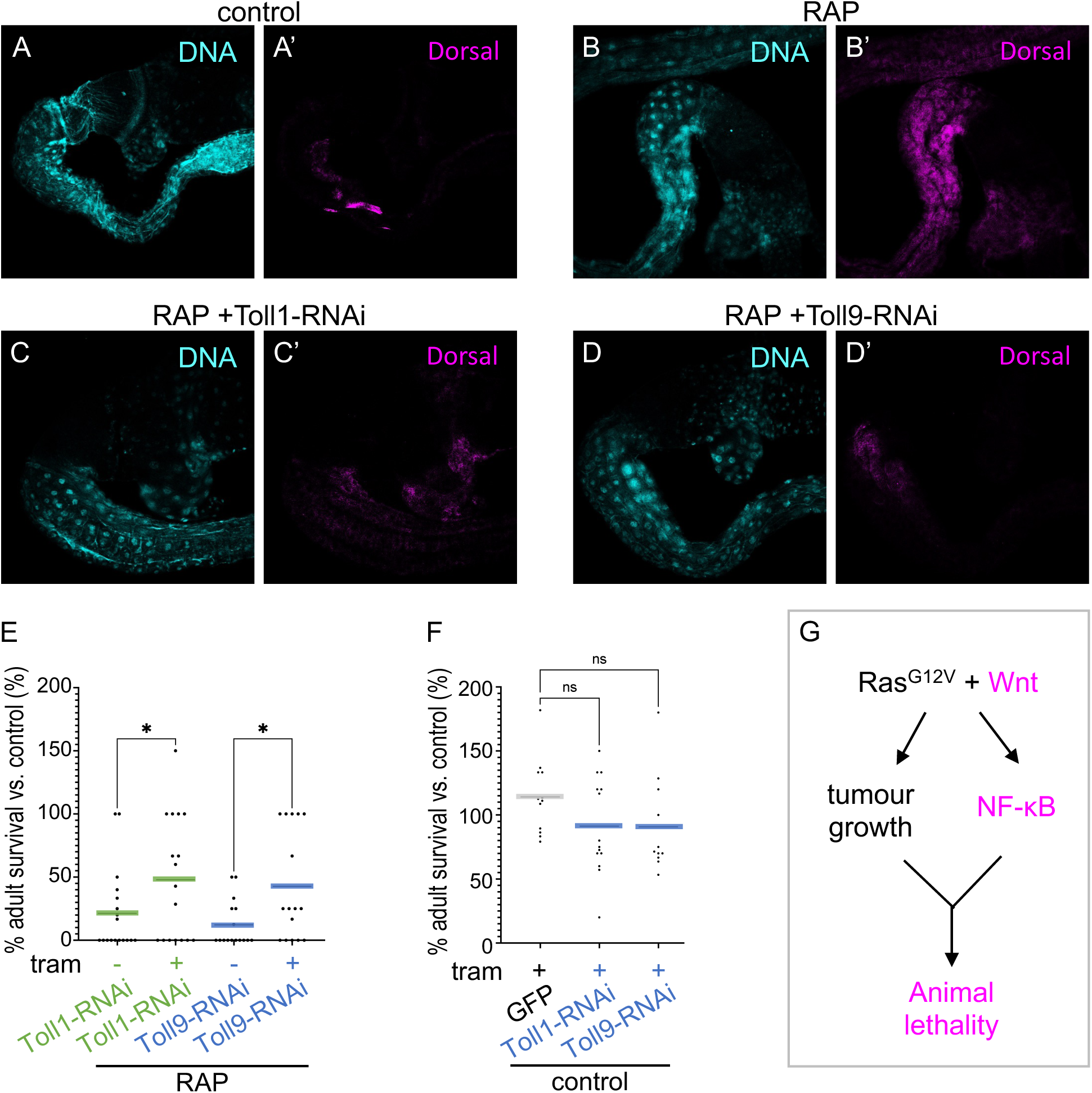
Toll-1 and Toll-9 are required for upregulation of canonical NF-κB activity in RAP tumours. **(A-D)** Control **(A)**, *RAP* **(B)**, *RAP* +*Toll1-RNAi* **(C)**, *RAP* +*Toll9-RNAi* **(D)** were induced in hindguts and were stained with anti-dorsal antibody. DNA are visualized with Propidium iodide (PI). **(E, F)** Percent survival of transgenic flies to adulthood relative to control flies was quantified in the present or absence of trametinib (1 μM). **€** *RAP* +*Toll1-RNAi* and *RAP* +*Toll9-RNAi*; **(F)** *Toll1-RNAi* and *Toll9-RNAi*. **(G)** A model for WNT activity and p53 defect promoting drug resistance in CRC.

These results suggest that Toll-1 and Toll-9 are required for enhancing canonical NF-κB activity in RAP tumours. Consistence with our previous results, knockdown of Toll-1 or Toll-9 did not significantly suppress tumour overgrowth of HZP in the presence of trametinib in *byn>RAP* tumours (Supplemental Figure 2a and S2b compared to Figure 1H, quantified in S1c). Reduction of *dl* in the whole animal setting using a loss of function mutation allele (*dl*[1]) significantly suppressed tumour-induced lethality in *byn>RAP* tumours in the presence of trametinib (Supplemental Figure 2g). However, administering NF-κB inhibitors such as QNZ (EVP4593) and JSH-23 does not strongly inhibit tumour-induced lethality in the presence of trametinib (Supplemental Figure 2d-f).

### WNT inhibitors increased trametinib efficacy on RAP tumours

Our results indicate that WNT activation plays a role in regulating host lethality by increasing canonical NF-κB activity and, in turn, opposing trametinib’s ability to reduce tumour overgrowth in *byn>Ras^G12V^* animals (Figure 2G). We therefore next assessed whether inhibiting WNT activity would reverse drug resistance in RAP tumours by feeding *byn>RAP* flies one of several (Figure 3A) WNT inhibitors plus trametinib. The result was emergent rescue: in particular, WNT pathway inhibitors PNU-74654 (*19*) and LF3 (*20*)—which suppress the interaction between β-Catenin and TCF—demonstrated strong efficacy in suppressing RAP tumours when paired with trametinib (Figure 3). This suppression led to increased animal survival in the presence of trametinib, without impacting survival in the absence of trametinib or in control animals (Figure 3A-3D, Supplemental Figure 3a-3d). Consistent with our results, combining trametinib plus PNU-74654 strongly suppressed canonical NF-κB activity in *byn>RAP* tumours (Supplemental Figure 3e).

**Figure 3:**
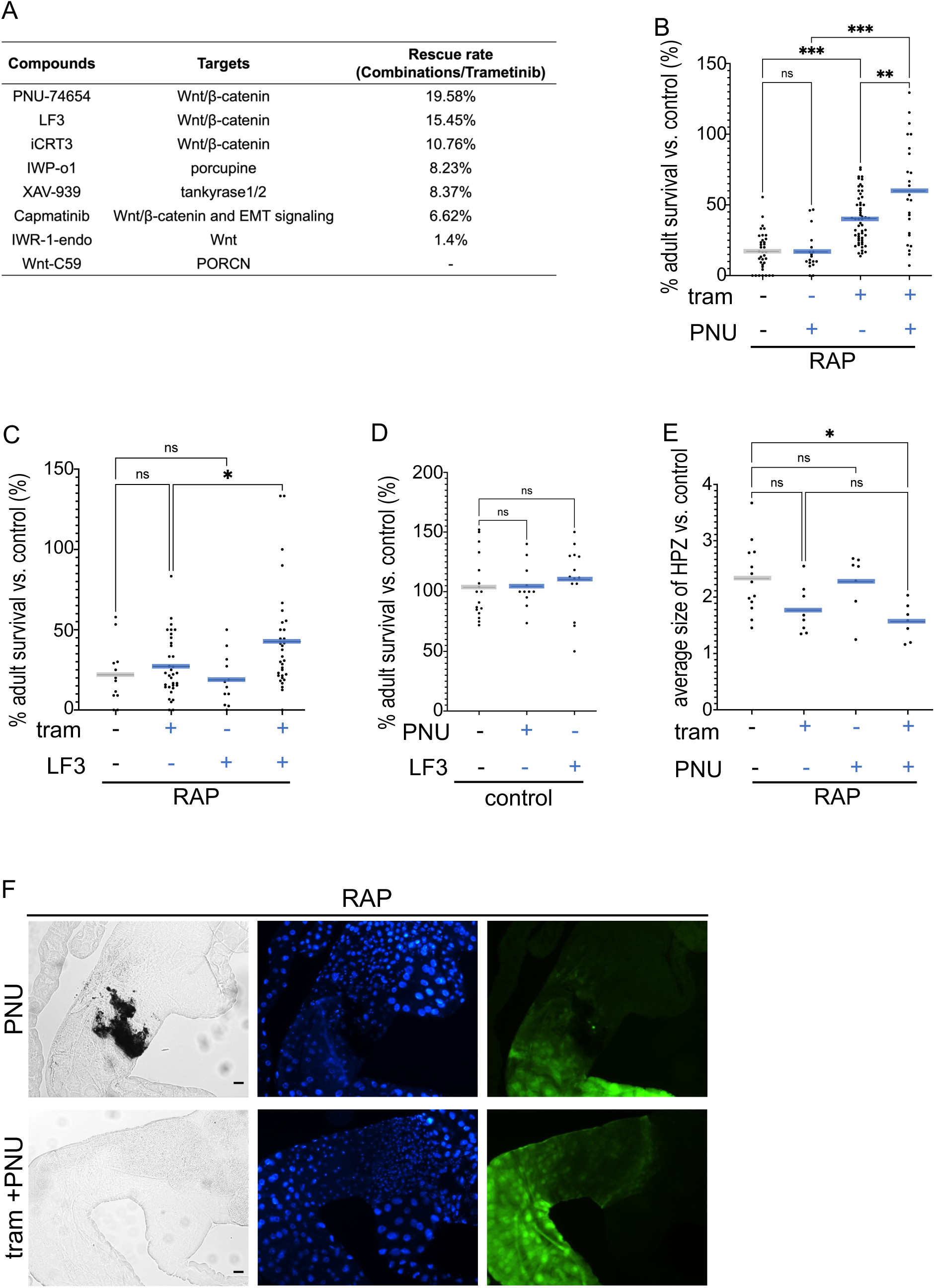
PNU-74654 and LF3 suppressed trametinib resistance in RAP tumours. **(A)** A summary of rescue rate of trametinib and WNT inhibitors drug combination in *RAP* hindgut tumours. **(B-D)** Percent survival of *byn>RAP* flies to adulthood relative to control flies was quantified in the present or absence of trametinib (1 μM), PNU-74654 (1 μM) or LF3 (10μM). **(E)** The average of hindgut proliferation zone (HPZ) size was measured by Fiji ImageJ and quantified as relative size to wild type (WT) hindgut. **(F, G)** Images of the digestive tract of third instar larvae in the present or absence of trametinib (1 μM) or PNU-74654 (1 μM).

Regarding tumour progression, PNU-74654 plus trametinib suppressed tumour overgrowth in the HPZ, while PNU-74654 alone did not affect tumour overgrowth (Figure 3E-3G, compared to Figure 1H). These data indicate that targeting WNT activity pharmacologically is effective at reducing trametinib resistance in RAP tumours.

### Combining trametinib and PNU-74654 suppressed tumour progression in genetically complex tumours

In previous work (*21*), we found that fly CRC models targeting 3-4 genes responded poorly to trametinib as a single agent, requiring drug combinations for efficacy. We therefore assessed whether combining trametinib plus PNU-74654 could effectively suppress tumour progression in still more genetically complex CRC lines. We tested seven ‘patient-specific fly avatar’ lines, each targeting 6-10 genes to more fully model the mutation profile of individual CRC patients (Supplemental Figure 4a). Each exhibited minimal response to trametinib alone (Figures 4, 5). In contrast, 5 of 7 avatar lines responded significantly to oral trametinib plus PNU-74654 (Figure 4A-4E), while two did not significantly respond to the cocktail (Figure 5I).

**Figure 4:**
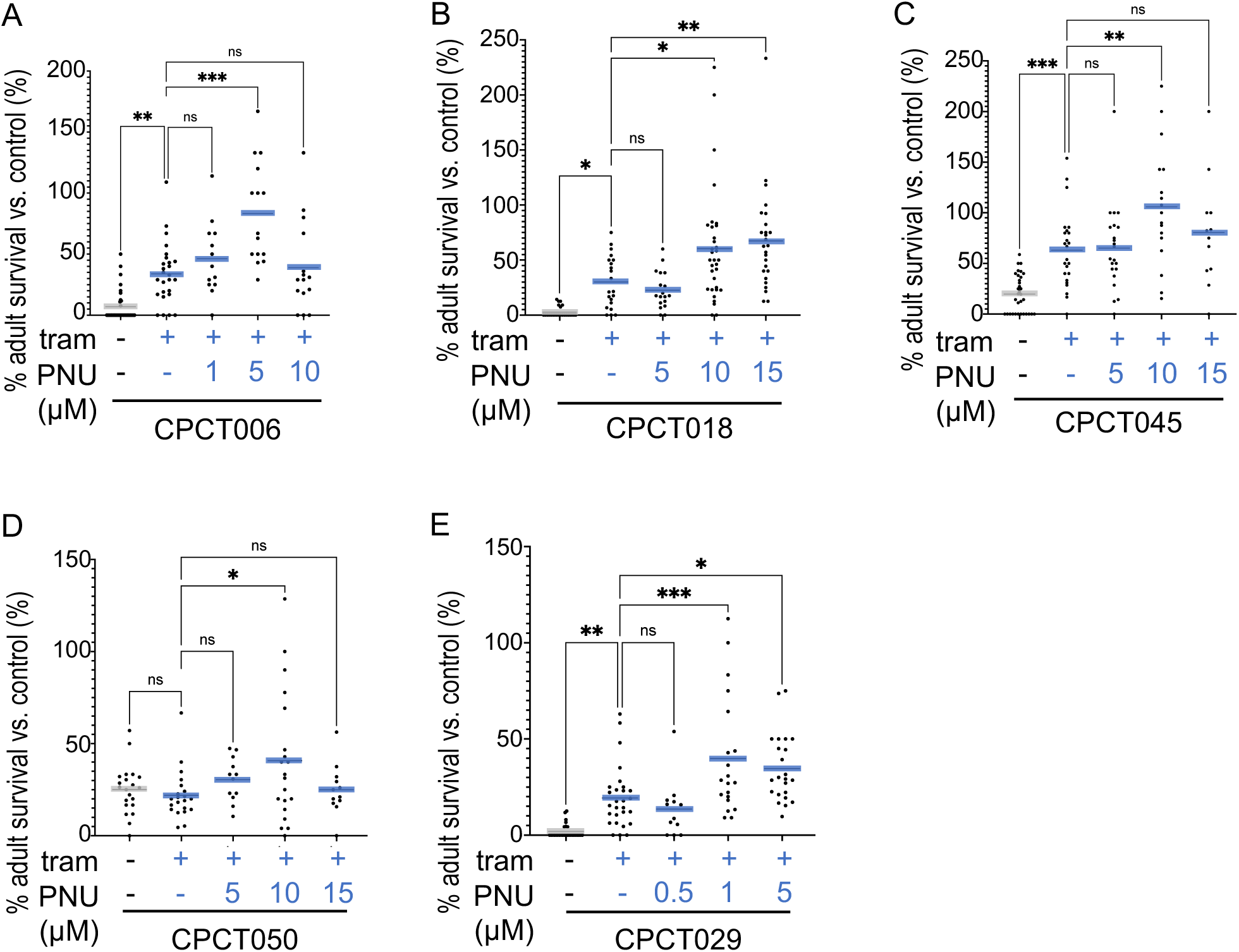
Combination of trametinib and PNU-74654 suppressed tumour progression in various genetically complex tumours. **(A-E)** Percent survival of transgenic patient-specific avatar fly lines to adulthood relative to control flies was quantified in the present or absence of trametinib (1 μM) or PNU-74654. **(A)** *CPCT006*; **(B)** *CPCT018*; **(C)** *CPCT045*; **(D)** *CPCT050*; **(E)** *CPCT029*.

**Figure 5:**
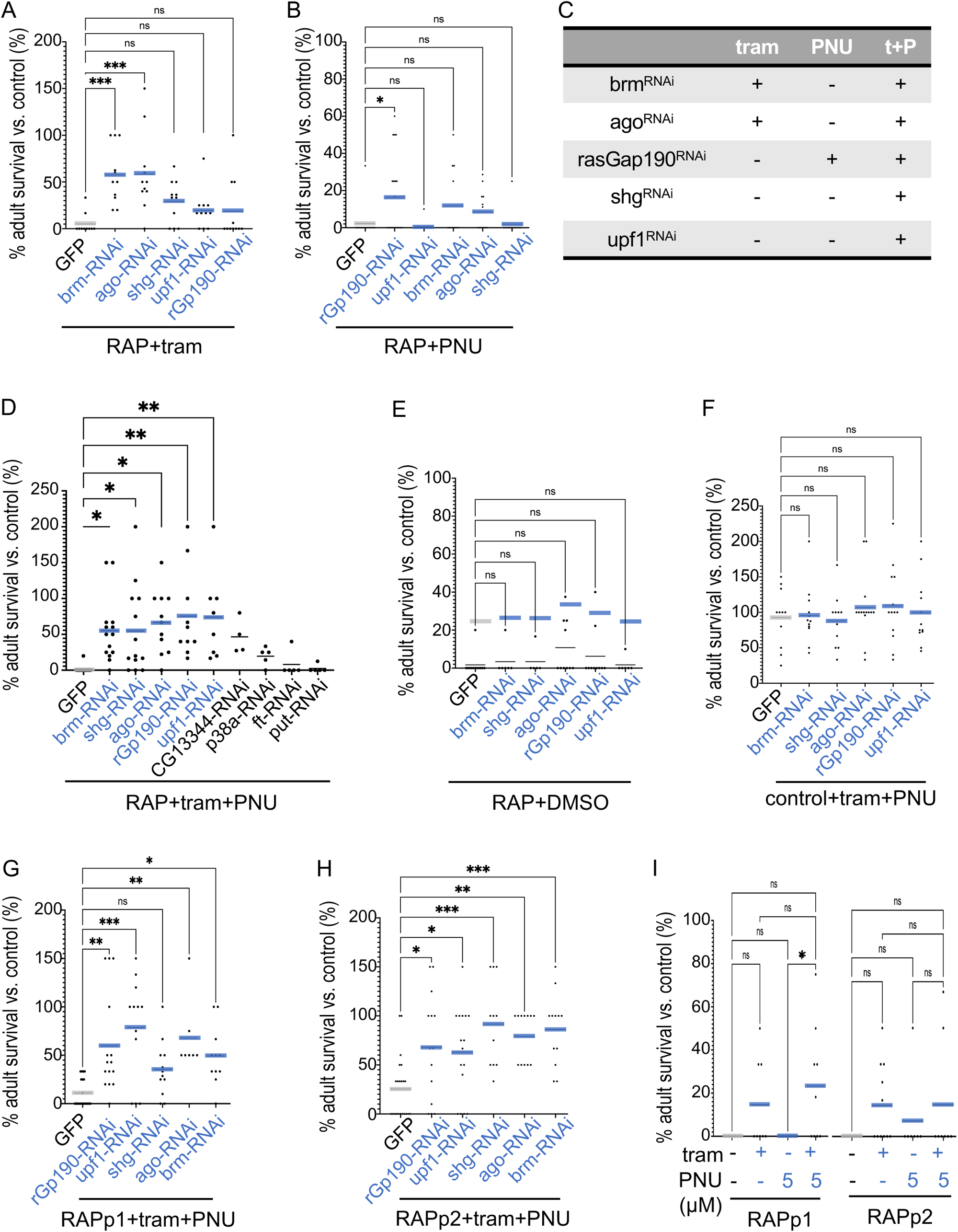
Regulators of combination of trametinib and PNU-74654 in CRC tumours. Survival of transgenic flies to adulthood relative to control flies was quantified in the present or absence of trametinib (1 μM) or PNU-74654 (1 µM except where noted). **(A)** RNAi-mediated knockdown of *brm* or *ago* improved survival of *byn>RAP* flies treated with trametinib. **(B)** *rhoGAPp190* knockdown improved survival of *RAP* flies treated with PNU-74654. **(C)** Summary of five loci found to impact *RAP* response to trametinib and/or PNU-74654. **(D)** Knockdown of *brm*, *shg*, *ago*, *rhoGAPp190*, or *upf1* improved survival to adulthood of *RAP* flies in the present of trametinib plus PNU-74654. These five loci did not significantly rescue *RAP* fly survival in the absence of drug **(E)**, nor did they alter survival of control animals in the presence of drug **(F)**. **(G, H)** Knockdown of *brm*, *shg*, *ago*, *rhoGAPp190*, or *upf1* rescued more genetically complex avatar lines RAPp1 and RAPp2 when treated with trametinib plus PNU-74654 (5 µM); note *shg* rescue of RAPp1 did not rise to the level of statistical significance. **(I)** In the absence of knockdown, trametinib and/or PNU-74654 (5 µM) exhibited poor rescue of RAPp1 and RAPp2.

### Regulators for combination of trametinib and PNU-74654 in CRC tumours

To identify genes that mediate the efficacy of trametinib plus PNU-7654 on RAP flies, we performed a genetic screen. We found five genes that, when targeted for knockdown by RNA-interference (RNAi), enhanced adult eclosion of *byn>RAP* flies in the presence of trametinib plus PNU-7654 (or LF3): *brahma* (*brm*, orthologue of human *SMARCA2* and *SMARCA4*), *shotgun* (*shg, CDH1*), *archipelago* (*ago*, *FBXW7*), *rhoGAPp190* (*rhoGAPp190*, *ARHGAP5, ARHGAP35*) and *upf1 RNA helicase* (*upf1*, *UPF1*). Importantly, none of these knockdowns impacted survival of control animals or untreated *byn>RAP* flies (Figure 5, Supplemental Figure 5). These data suggest that *brm*, *shg*, *ago*, *rhoGAPp190* and *upf1* help mediate efficacy of trametinib plus a WNT pathway inhibitor in CRC tumours. Testing single drugs with each of the five loci, we found that *brm-RNAi* and *ago-RNAi* significantly increased the sensitivity of *byn>RAP* tumours to trametinib (Figure 5A, 5C); *rhoGAPp190-RNAi* sensitized *byn>RAP* tumours to PNU-74654 (Figure 5B, quantified in 5C).

As noted above, 2 of 7 tested ‘patient-specific fly avatar’ lines, RAPp1 and RAPp2, were resistant to combined trametinib plus PNU-7654 (Figure 5I). Knockdown of *brm*, *shg*, *ago*, *rhoGAPp190* and *upf1* each enhanced tumour sensitivity to trametinib plus PNU-7654 in both multigenic tumours with the exception that *shg* had only a weak effect on *byn>RAPp1* drug response (Figure 5F and 5G). Finally, we note human orthologs of these five loci are altered in a subset of human CRC patients (Supplemental Figure 5a), suggesting they could serve as biomarkers for patients that would be especially sensitive to trametinib plus PNU-74654.

### Elevated WNT plus KRAS is associated with increased canonical NF-κB signalling in human CRC tumour samples

To assess if our Drosophila data is relevant to human CRC, we examined human CRC tissue sections to determine whether high WNT activity is associated with high NF-κB activity in samples with oncogenic KRAS mutations; we used IKK isoforms to identify canonical vs. non-canonical NF-κB signalling. IKKβ serves as the primary catalytic subunit of IKK, activating canonical NF-κB signalling by proinflammatory cytokines such as TNFα, IL-1, and LPS. In contrast, IKKα activates non-canonical NF-κB signalling, activated by other members of the TNFR superfamily (*22*, *23*): for example, IKKα is phosphorylated by NIK at Ser-176 to promote release of the non-canonical NF-κB factor RelB-p52 into the nucleus to activate target genes (*24*).

WNT activity was assessed in human CRC patient samples by anti-β-catenin antibody (Supplementary Figures 6a). We observed that the expression of the IKKβ protein was significantly higher in CRC patient samples with both high WNT activity *and* oncogenic KRAS compared to those with only high WNT activity *or* KRAS mutations (Figure 6A-D, 6I). In contrast, we found no significant differences in the levels of IKKα or Ser-176 phosphorylated IKKα in samples with (i) high WNT/oncogenic KRAS samples vs. (ii) high WNT activity or oncogenic KRAS (Figure 6E-H, 6J; Supplemental Figure 6b-f). These results are consistent with our fly data: in KRAS mutant CRC tumours, high WNT activity is associated with enhanced canonical NF-κB signalling.

**Figure 6:**
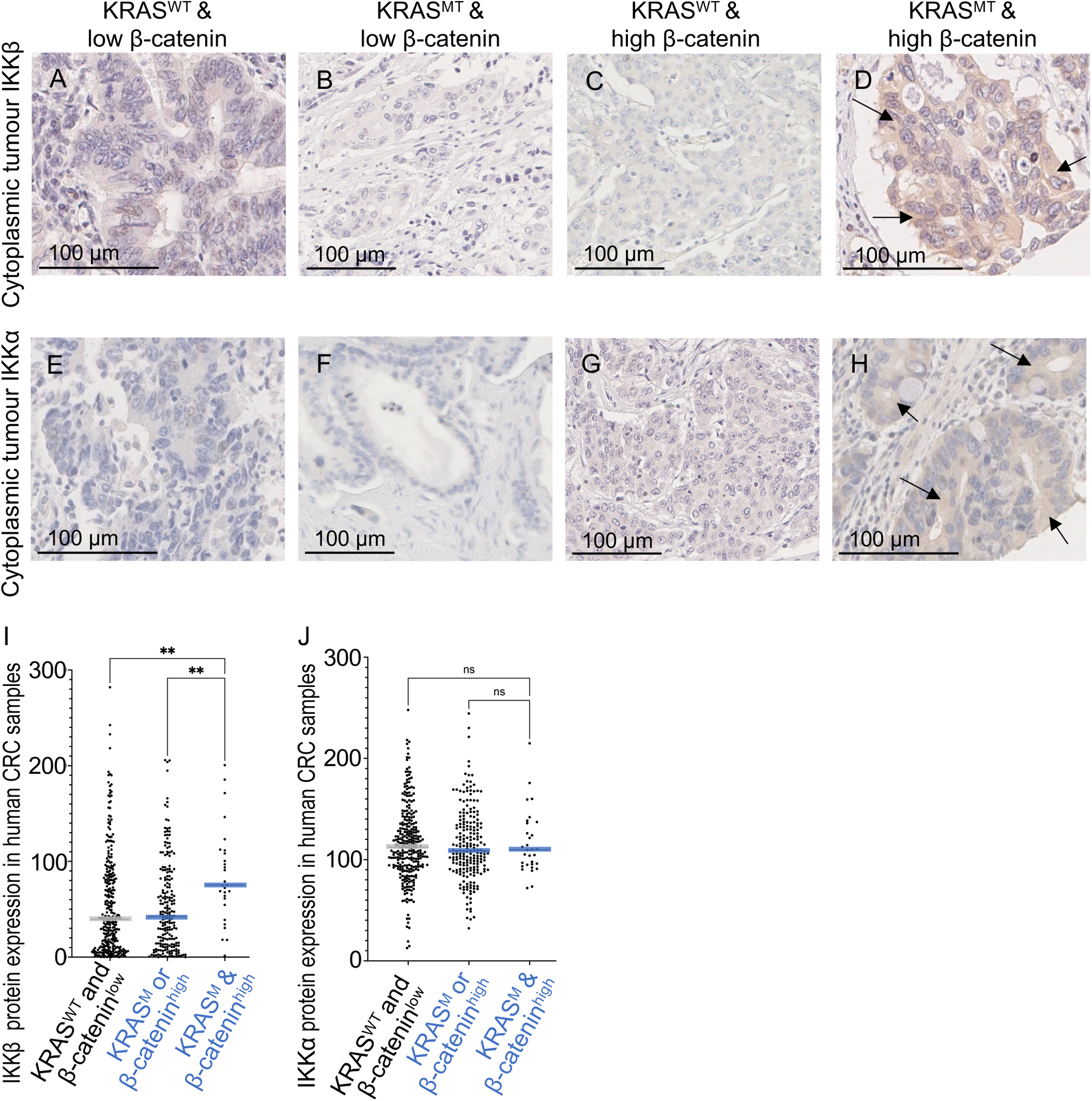
Co-activation of WNT and KRAS was associated with an upregulation of canonical NF-κB activity in human CRC. **(A-H)** Representative immunohistochemical (IHC) staining for IKKβ, IKKα in stage 2-3 colorectal cancer patient samples. **(A, E)** *KRAS^WT^* (wild-type) plus low β-catenin expression CRC samples; **(B, F)** *KRAS^MT^* (mutation in position G12/G13) plus low β-catenin expression CRC samples; **(C, G)** *KRAS^WT^*plus high β-catenin expression CRC samples; **(D, H)** *KRAS^MT^*plus high β-catenin expression CRC samples. **(I, J)** Graph shows the median of expression of IKKβ **(I)** or IKKα **(J)** in each different mutated human CRC, determined by IHC intensity values. Patients were grouped into 3 categories based on KRAS status and β-catenin expression.

## Discussion

In this study, we conduct an in-depth analysis of tumours that contain three genes commonly mutated in human CRC: *RAS* (typically *KRAS*), *APC,* and *P53* (‘RAP’). We found that elevated WNT activity led to upregulation of canonical NF-κB signalling when paired with RAS^G12V^ in *Drosophila* hindgut tumours, as well as in human tumours that paired elevated WNT activity with oncogenic KRAS isoforms. Elevated canonical NF-κB activity in turn reduced efficacy of the MEK inhibitor trametinib in RAP tumours. This resistance to trametinib was reversed when WNT inhibitors were included. Oncogenic RAS isoforms are present in approximately half of all CRC tumours, and current second-line treatments have shown limited efficacy.

Most WNT pathway inhibitors currently undergoing clinical trials act by promoting degradation of β-catenin, including inhibitors of PORCN (*e.g.,* ETC-1922159, WNT974 and XNW7201) and Frizzled receptors (*e.g.,* vantictumab, ipafricept) (*25*). However, our data suggests that WNT inhibitors such as PNU-74654 or LF3—which disrupt the interaction between β-catenin and TCF—are particularly effective when used in combination with trametinib for treating CRC: this drug combination proved effective even in more genetically complex CRC avatar lines. To extend our drug resistance work, we identified five genes that further enhanced efficacy of the drug combination: *brm*, *shg*, *ago*, *rhoGAPp190* and *upf1.* These genes are mutated in a subset of patients, identifying a cohort that may prove especially responsive to trametinib/WNT inhibitor drug combination. Two genes—*brm* and *ago*—enhanced the efficacy of trametinib alone, identifying a candidate biomarker for trametinib response. That is, our work identifies a path to matching drugs to specific subsets of RAS-mutant CRC patients, an especially challenging cohort.

The NF-κB pathway has been widely linked to cancer, including impacting drug resistance by regulating the survival of cancer cells. For example, NF-κB activity is reported to inhibit the response of HTC15 human colon cancer cells to daunomycin by controlling drug uptake (*13*). NF-κB activity is also linked to sorafenib resistance in CD13+ hepatocellular carcinoma cell lines by controlling genes that regulate cell cycle and apoptosis (*26*). Pharmacologically blocking the NF-κB pathway sensitizes tumour cells to doxorubicin in Dll1+ mouse breast cancer cells by promoting cell death (*27*). In this whole animal study, we demonstrate that canonical NF-κB-mediated drug resistance is an emergent property of CRC tumours that combine high WNT activity with oncogenic RAS. This may have therapeutic implications, as we demonstrate.

We previously showed a role for TNF signalling in regulating tumour progression in a RAS-dependent Drosophila cancer model (*28*) and, indeed, removing just one genomic copy of the *dl* gene was sufficient to significantly suppress tumour-induced animal lethality. Interestingly, pharmacological inhibition of canonical NF-κB pathway activity by JSH23 weakly *suppressed* RAP tumour-induced lethality while the NF-κB signalling inhibitor QNZ (EVP4593) *enhanced* trametinib resistance in RAP tumours. QNZ targets TNFα production, suggesting that systemic TNFα production plays a role in inhibiting tumour progression in the present of trametinib. Indeed, it has been reported that TNFα renders tumour vessels more permeable, facilitating the delivery of anticancer drug agents to solid tumours (*29*). These data offer guidance for development of a next generation of NF-κB inhibitors.

Genetic complexity is a common clinical feature of tumours. Here we link one version of this complexity linked to aggressive CRC disease—RAS, WNT, P53—as sufficient to direct overgrowth of the hindgut proliferative zone (HPZ) and promote emergent drug resistance. Currently, patient tumours with these three altered genes have few second line therapeutic options, as RAS pathway inhibitors have failed to provide durable regression. Gaining a deeper understanding of how this combination directs drug resistance through NF-κB provides a new candidate avenue towards therapeutics.

## Materials and methods

### *Drosophila* strains and genetics

Fly lines were cultured at room temperature or 25-29 °C on standard fly food or food-plus-compound. Fly food contained tayo agar 10g, soya flour 5g, sucrose 15g, glucose 33g, maize meal 15g, wheat germ 10g, treacle molasses 30g, yeast 35g, nipagin 10ml, propionic acid 5ml in 1000 ml water. Transgenes used (Bloomington Drosophila Stock Center number): *byn-gal4* (hindgut-specific line, V. Hartenstein), *UAS-Ras^G12V^* (second chromosome, G. Halder), *tub-gal80^TS^* (#7017), *w^1118^* (#3605), *UAS-mCD8-GFP* (#5137), *UAS-dl-RNAi* (#36650), *UAS-dif-RNAi* (#30513), *UAS-cact-RNAi* (#37484), UAS-Arm^S10^ (#4782), *UAS-Toll1-RNAi* (#35628), *UAS-Toll9-RNAi* (#34853), *UAS-brm-RNAi* (#35211), *UAS-shg-RNAi* (#38207), *UAS-ago-RNAi* (#34802), *UAS-rhoGAPp190-RNAi* (#43987), *UAS-upf1-RNAi* (#64519), *UAS-CG13344-RNAi* (#41831), *UAS-p38a-RNAi* (#35244), *UAS-ft-RNAi* (#34970), *UAS-put-RNAi* (#39025), *UAS-dnapol-eta-RNAi* (#33410), *UAS-lrp1-RNAi* (#44579), *UAS-tefu-RNAi* (#44073), *UAS-nej-RNAi* (#37489), *UAS-nos-RNAi* (#33973), *UAS-pc-RNAi* (#36070), *UAS-rad51c-RNAi* (#67355), *dl[1]* (#3236).

### Chemicals

Drugs and compounds were used as follows: trametinib (Selleckchem or biorbyt), QNZ (EVP4593), iCRT3, IWP-01, XAV-939, Capmatinib, JSH-23, PNU-74654 and Propidium iodide (PI) purchased from Selleckchem. Drug and compound stocks were diluted in DMSO or water; drugs were then mixed into standard fly food with final DMSO concentration 0.1% to prevent toxicity.

### Statistical analysis (*Drosophila*)

Eggs were collected for 24 hours in drug-containing food at 18 °C to minimize transgene expression during embryogenesis to prevent embryonic effects or lethality. After 3 days, the tubes were transferred to the appropriate temperature to induce transgene expression; the number of surviving Drosophila adults was quantified after 2 weeks. *byn>w^1118^* served as a control in this study. Statistical analysis was performed using Prims9. N.S P(>0.12), * P(0.033), ** P(0.002), *** P(0.001), and **** P(<0.0001). All statistical data are summarized in Supplementary Table 1. All detailed genotypes are summarized in Supplementary Table 2.

### Imaging of the digestive tract of third instar larvae

Third instar larvae were dissected in 1x PBS and fixed with 4% paraformaldehyde for 30 min at room temperature, then washed 3×15 min in PBT (0.1% Triton X in 1x PBS). Samples were incubated in anti-dorsal primary antibody (#7A4, DSHB, 1:100); the secondary antibody used was anti-mouse Alexa Fluor 546 or 647 (Invitrogen, 1:250). Samples were mounted with DAPI-containing SlowFade Gold Antifade Reagent (#S36939, Molecular Probes). Fluorescence images were visualized on a Lecia TSC SPE confocal microscope.

### RNA isolation and quantitative real time PCR (*Drosophila*)

Total RNA from 30 hindguts was isolated using TRIzol® according to the manufacturer’s protocol (cat.15596018, Invitrogen™, Life Technologies). mRNA was reverse transcribed using iScriptTM gDNA Clear cDNA Synthesis Kit (cat# 1725035, Bio-Rad Laboratories Ltd).

For quantitative Real-Time PCR (qPCR), iTaq™ Universal SYBR® Green Supermix kit (cat. #1725124, Bio-Rad Laboratories Ltd.) was used according to the manufacturer’s recommendation with cDNA (diluted 1:10∼20) as a template. RT-qPCRs were performed with three biological replicates. Relative expression values were determined by the 2^−ΔΔCt^ method using *rp49* as endogenous control. The RT-qPCRs primers used as following: *rp49* (forward: CGCTTCAAGGGACAGTATCTG; reverse: AAACGCGGTTCTGCATGA), *drosomycin* (forward: CTCTTCGCTGTCCTGATGCT; reverse: ACAGGTCTCGTTGTCCCAGA).

### Immunohistochemistry for detection of β-catenin, IKKβ, IKKα and phospho-IKKα^s176^

Samples from a retrospective cohort of 787 stage 2-3 colorectal cancer patients were stained via immunohistochemistry (IHC) for β-catenin, IKKβ, IKKα and phospho-IKKα serine 176 (IKKα^s176^). Staining was performed on a previously constructed tissue microarray (TMA), which consisted of CRC tissue from patients undergoing surgery with curative intent within Greater Glasgow and Clyde hospitals between 1997-2013. Data are stored within the Glasgow Safehaven (GSH21ON009) and ethical approval was in place for the study (MREC/01/0/36).

IHC was performed as previously described (*30*). Briefly, TMA sections were dewaxed then rehydrated through a serious of alcohols. Antigen retrieval was performed using citrate buffer (pH6) for β-catenin, IKKβ and IKKα, and Tris EDTA (pH 9) for IKKα^s176^. Endogenous peroxidases were blocked in 3% hydrogen peroxide. Tissue was blocked using 10 % casein (SP-5020, Vector laboratories, CA, USA) for β-catenin and IKKα^s176^, and 5% horse serum (S-2000, Vector laboratories, CA, USA) for IKKβ and IKKα, incubating for 1 hour at room temperature. Sections were incubated in primary antibody β-catenin ((M3539, Dako, CA, USA, 1:600), IKKα (GWB-662250, Genway, CA, USA, 1:4000), IKKβ (ab32135, abcam, Cambridge, UK, 1:200) and IKKα^s176^(ab138426, abcam, Cambridge, UK, 1:150) overnight at 4°C. Sections were washed in tris-buffered saline (TBS), incubated in Impress secondary antibody (MP-7500, Vector laboratories, CA, USA) for 2 hours at room temperature. Sections were washed in TBS and incubated for 5 minutes in 3,3′-Diaminobenzidine (DAB) (SK-4105, Vector laboratories, CA, USA). Slides were rinsed in water, counterstained and dehydrated before mounting with Pertex (00801-EX, Histolab products, Askim, Sweden). Stained sections were imaged using a Hamamatsu NanoZoomer (Hamamatsu Photonics, Shizuoko, Japan) onto NZ Connect viewing platform (Hamamatsu Photonics, Shizuoko, Japan).

### Staining quantification of β-catenin, IKKβ, IKKα and phospho-IKKα^s176^ (human CRC)

Staining intensity was assessed semi-quantitively by weighted histoscore using QuPath® software in the tumour cell cytoplasm for β-catenin, IKKβ, IKKα and IKKα^s176^ (*31*). Continuous scores ranging from 0-300 for β-catenin were dichotomised into high and low expression groups using the Survminer package in RStudio (version 1.4, RStudio, Boston, MA, USA).

### Mutational profiling and Analysis (human CRC samples)

CRC tissue from the patient cohort was profiled for presence of KRAS mutation by BioClavis (BioClavis Ltd, Glasgow, UK). Patients were grouped into three categories based on KRAS status and β-catenin expression. Group 1 patients were wild type for KRAS and low for β-catenin, Group 2 patients were either KRAS mutant or high for β-catenin, and Group 3 patients were both KRAS mutant and high for β-catenin. These groups were then assessed for association with IKKβ, IKKα and IKKα^s176^ expression using T-tests in GraphPad Prism (GraphPad Software, La Jolla, CA, USA).

## Acknowledgements

We thank the Cagan Laboratory members for important discussions. We also thank the Bloomington Drosophila Stock Center for *Drosophila* stocks. This work was generously supported by grants from the NIH (R01CA258736) and a Royal Society Wolfson Fellowship. Some sentences were revised by GPT-3.5 for clarity.

## Compliance with ethical standards

The authors declare that they have no conflict of interest.

## Data and materials availability

The paper and/or Supplementary Materials contain all the necessary data for assessing the draw conclusions.

## Supporting Information

**Supplementary Figure 1:**
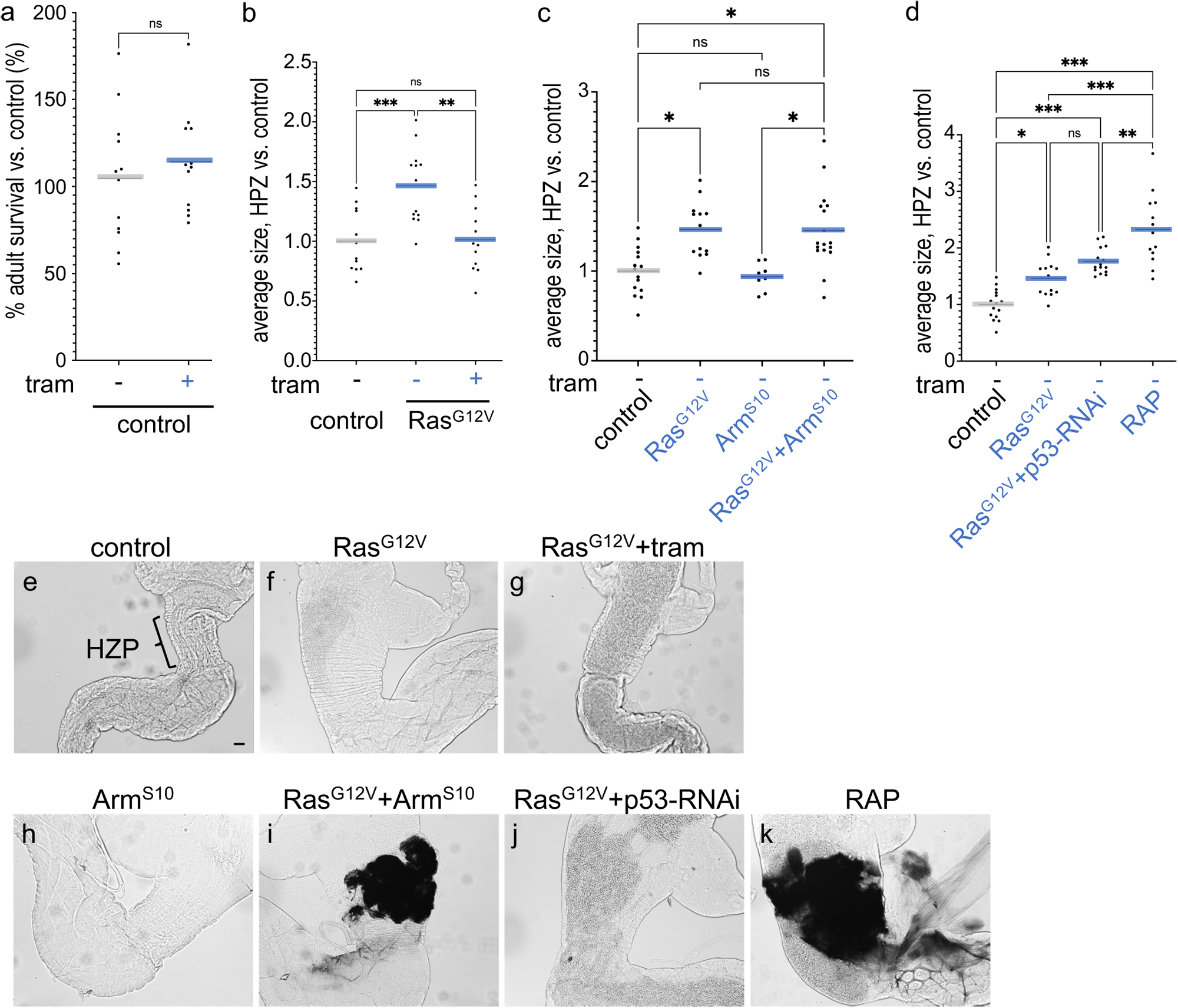
Overgrowth of HPZ was driven by Ras, Wg activity and p53 defect. (a) Percent survival of control flies to adulthood relative to control flies was quantified in the present or absence of trametinib (1 μM). (b-d) The average of hindgut proliferation zone (HPZ) size was measured by Fiji ImageJ and quantified as relative size to control hindgut. (e-k) Images of the digestive tract of third instar larvae in the present or absence of trametinib (1 μM). control (e), *Ras^G12V^* (f and g), *Arm^S10^* (h), *Ras^G12V^* +*Arm^S10^* (i), *Ras^G12V^* +*p53-RNAi* (j) and *RAP* (k).

**Supplementary Figure 2:**
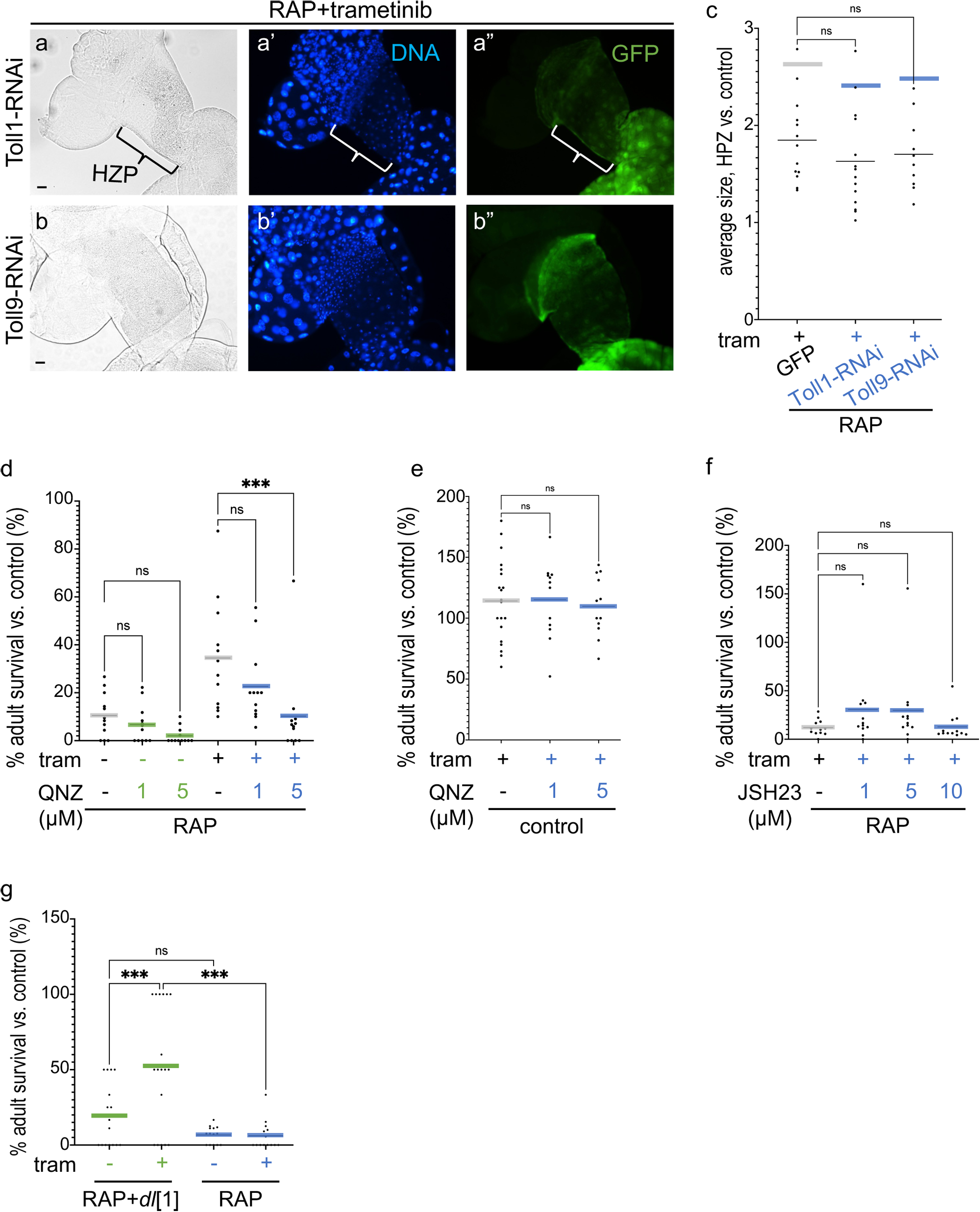
Administration of NF-κB inhibitors did not suppress drug resistance in RAP tumours. (a and b) Images of the digestive tract of third instar larvae in the present of trametinib (1 μM). (c) The average of hindgut proliferation zone (HPZ) size was measured by Fiji ImageJ and quantified as relative size to control hindgut. (d-g) Percent survival of transgenic flies to adulthood relative to control flies was quantified in the present or absence of trametinib (1 μM), QNZ (EVP4593) or JSH23. (d and f) *RAP*; (e) control; (g) *RAP*+ *dl[1]* and *RAP*.

**Supplementary Figure 3:**
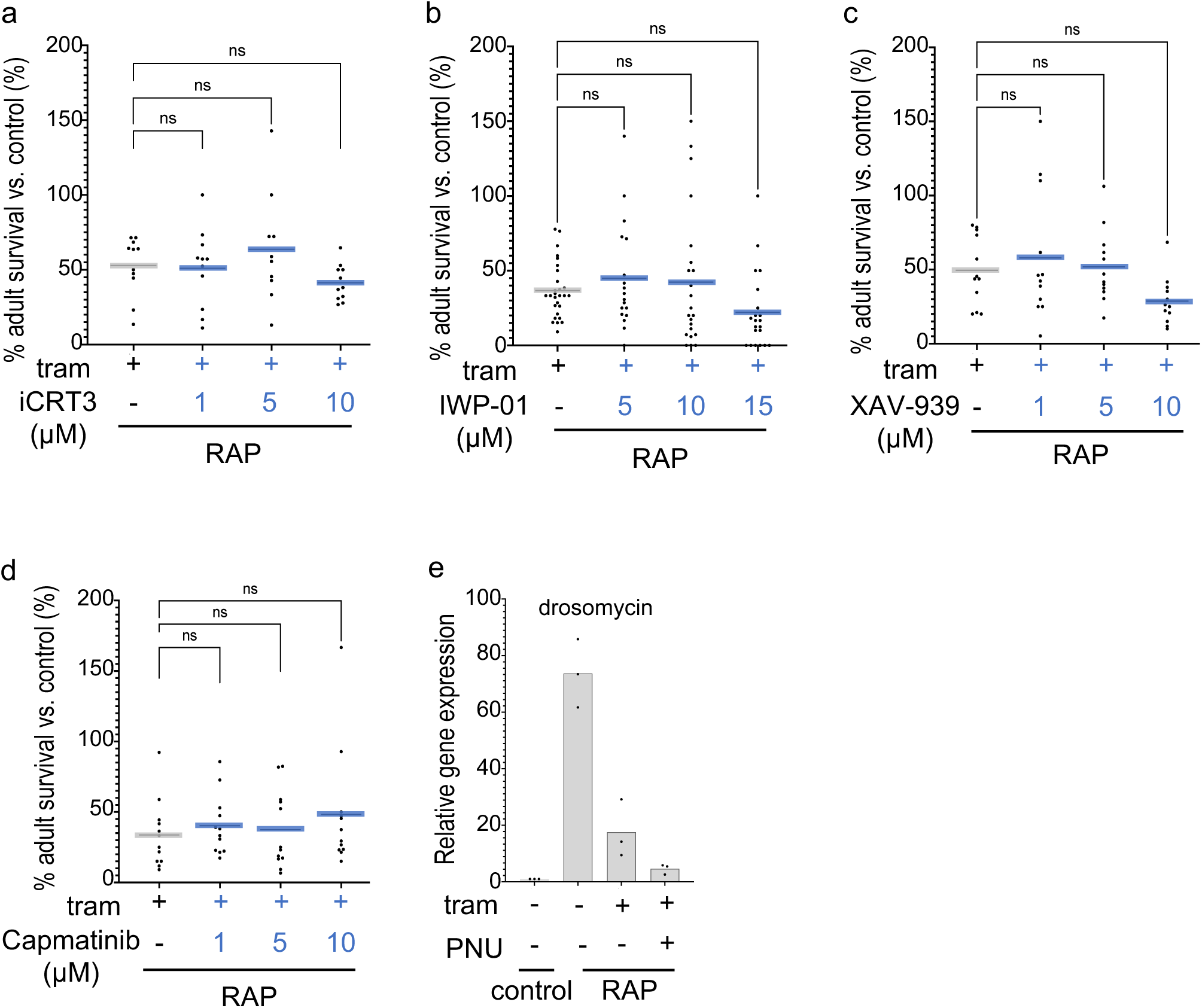
A screening for trametinib and WNT inhibitors drug combination in RAP hindgut tumours. (a-d) Percent survival of transgenic *RAP* flies to adulthood relative to control flies was quantified in the present or absence of trametinib (1 μM), iCRT3, IWP-01, XAV-939 or Capmatinib. (e) The expression of levels of drosomycin among each genotype in the present or absence of trametinib (1 μM) or PNU-74654 (1 μM) were detected by quantitative RT-PCR. (e) control and *RAP*.

**Supplementary Figure 4:**
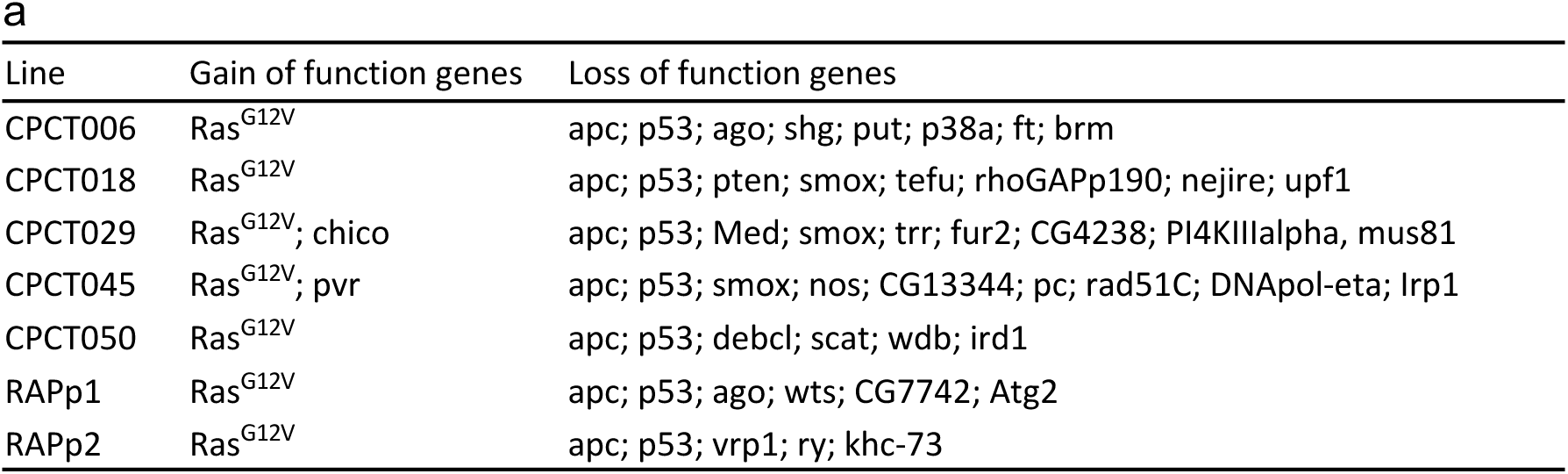
Mutations in CRC models. (a) A summary of mutations in patient-specific CRC models. These patient-specific fly avatars come from a fly-to-bedside study (CPCTs). and TCGA (p1, p2).

**Supplementary Figure 5:**
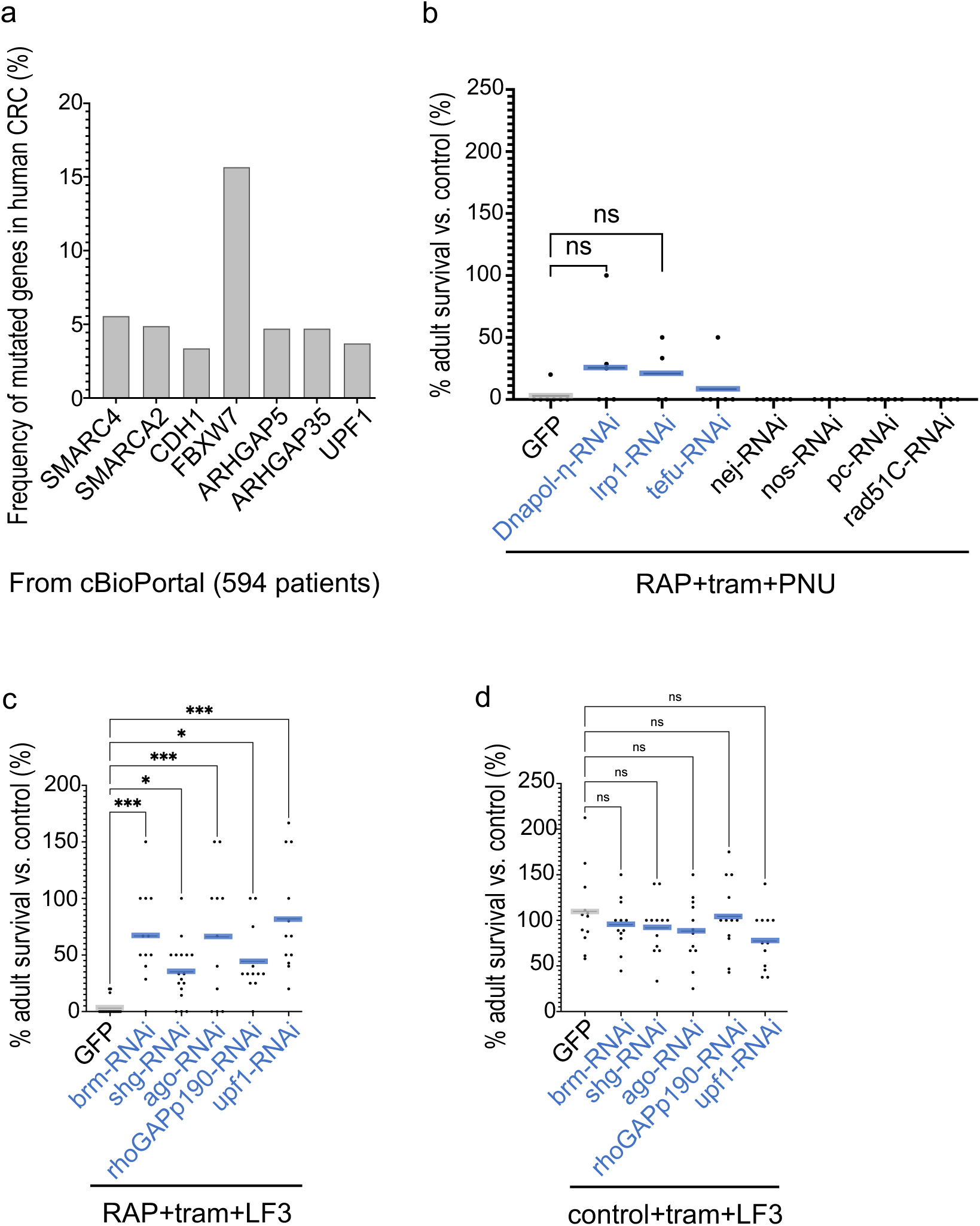
Regulators for combination of trametinib and LF3 in CRC tumours. (a) Graph showed frequency of mutated genes in human CRC. These data from cBioPortal including 594 patients. (b-d) Percent survival of transgenic flies to adulthood relative to control flies was quantified in the present or absence of trametinib (1 μM), PNU-74654 (1 μM) or LF3 (10μM). (b and c) *RAP*; (d) control.

**Supplementary Figure 6:**
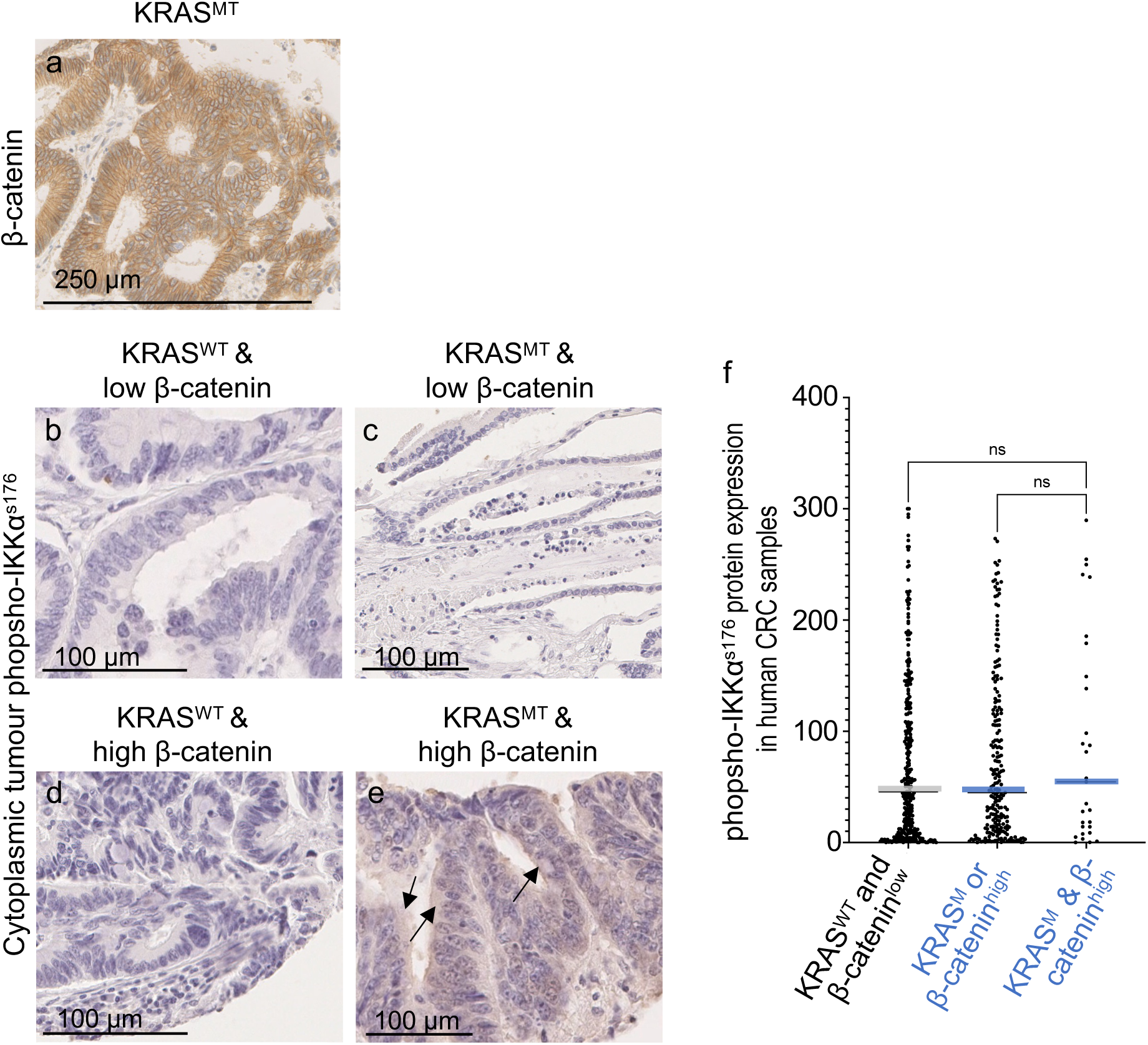
IHC staining of phospho-IKKα^s176^ protein in human CRC. (a-e) IHC staining of β-catenin or phospho-IKKα^s176^ in each genotype CRC sample. (a) *KRAS^MT^* (mutation in position G12/G13); (b) *KRAS^WT^* plus low β-catenin; (c) *KRAS^MT^* plus low β-catenin; (d) *KRAS^WT^* plus high β-catenin; (e) *KRAS^MT^* plus high β-catenin expression CRC samples. (f) Graph shows the median of expression of phospho-IKKα^s176^ in each different mutated human CRC, determined by IHC intensity values. Patients were grouped into 3 categories based on KRAS status and β-catenin expression.

**Supplementary Table1: a summary of statistics.**

**Supplementary Table2: a summary of detailed drosophila genotypes.**

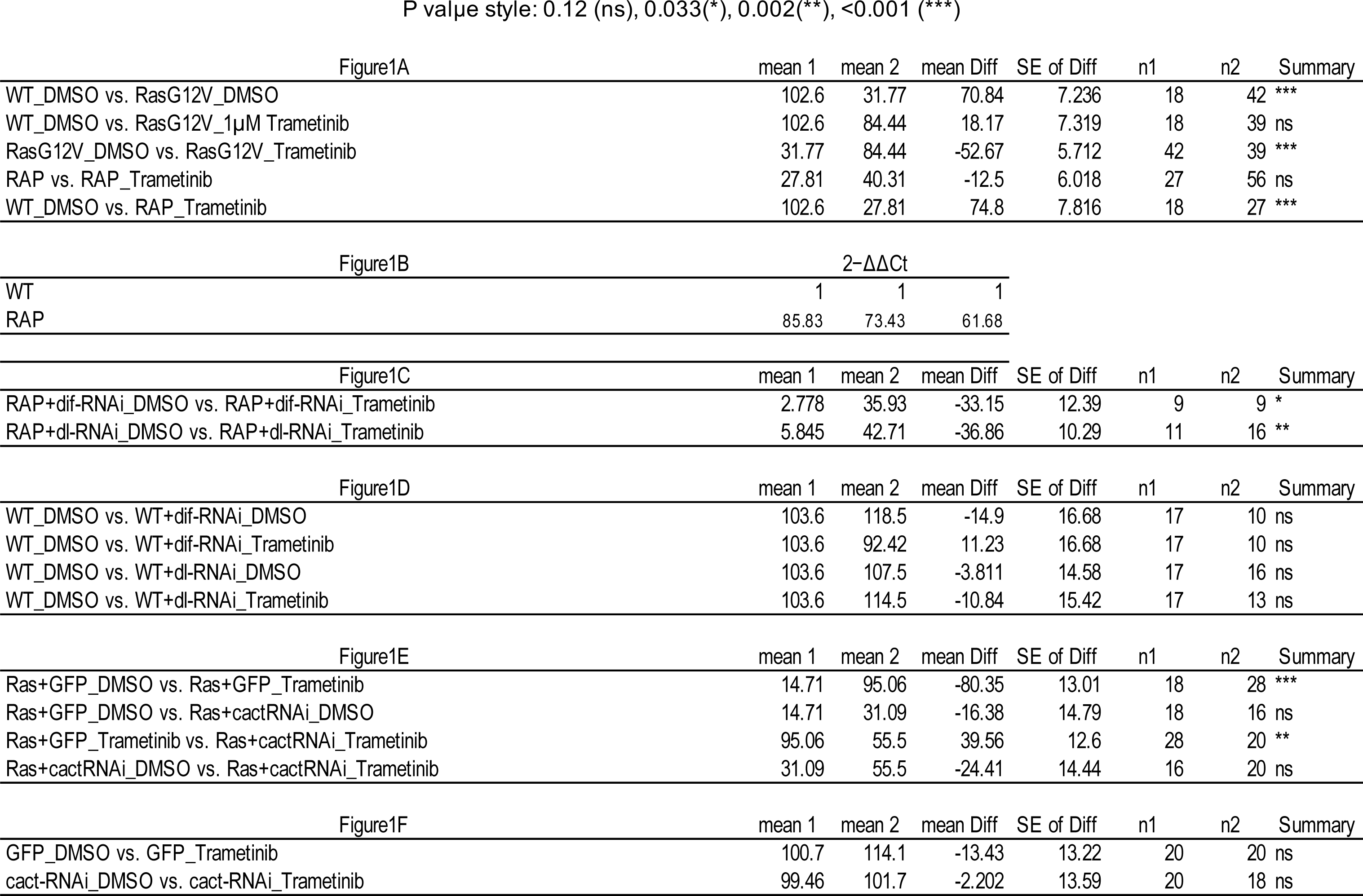
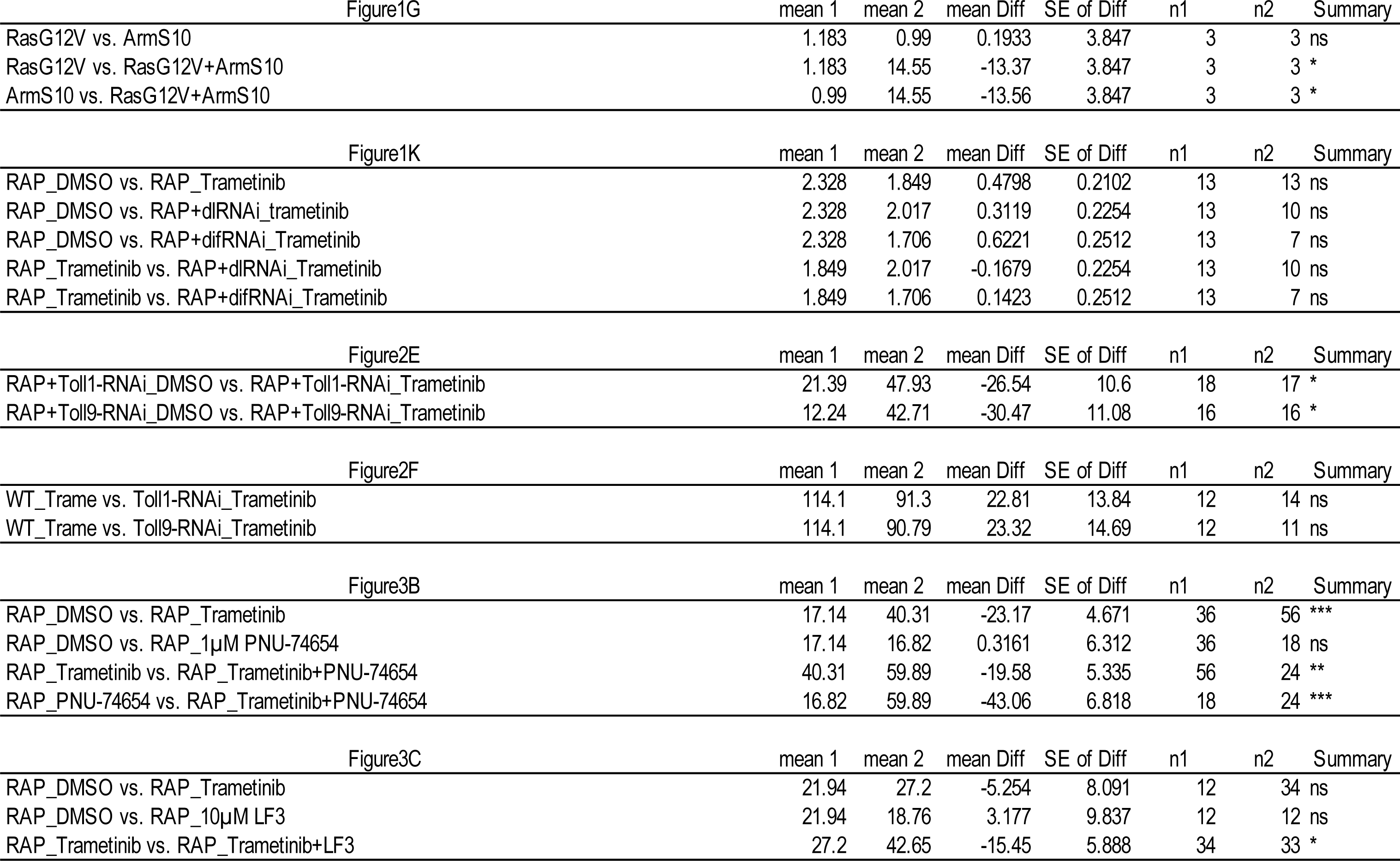
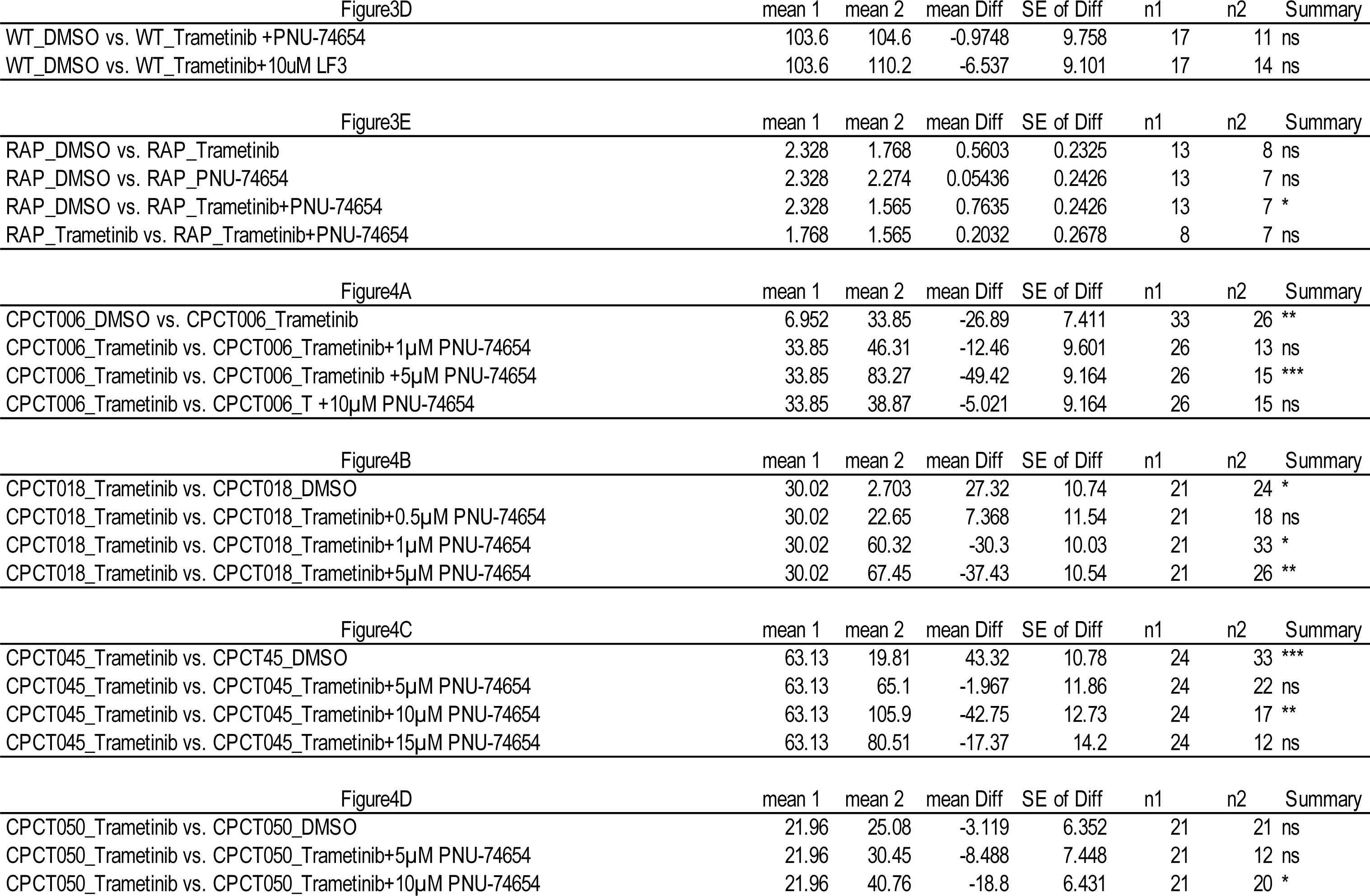
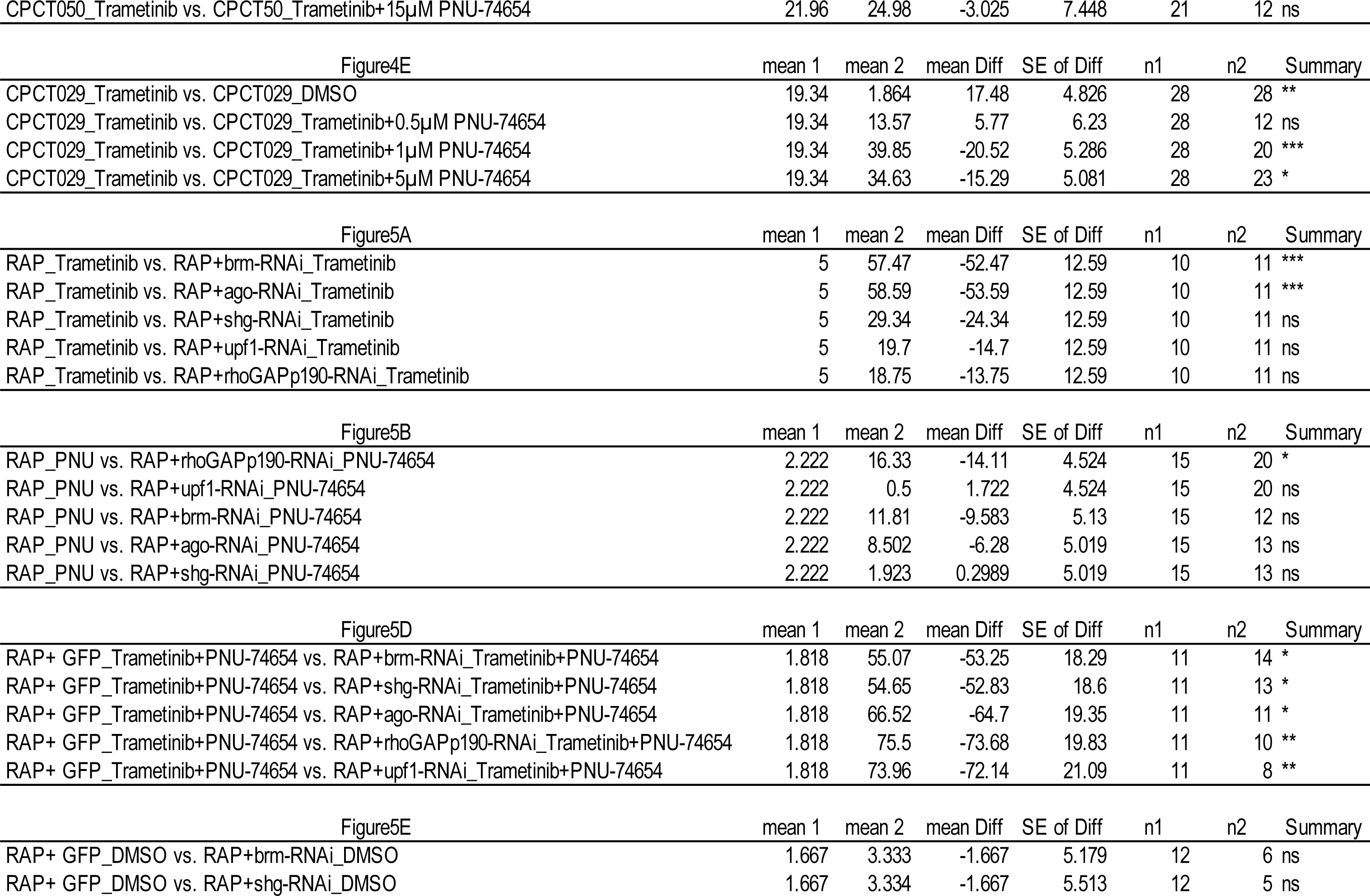
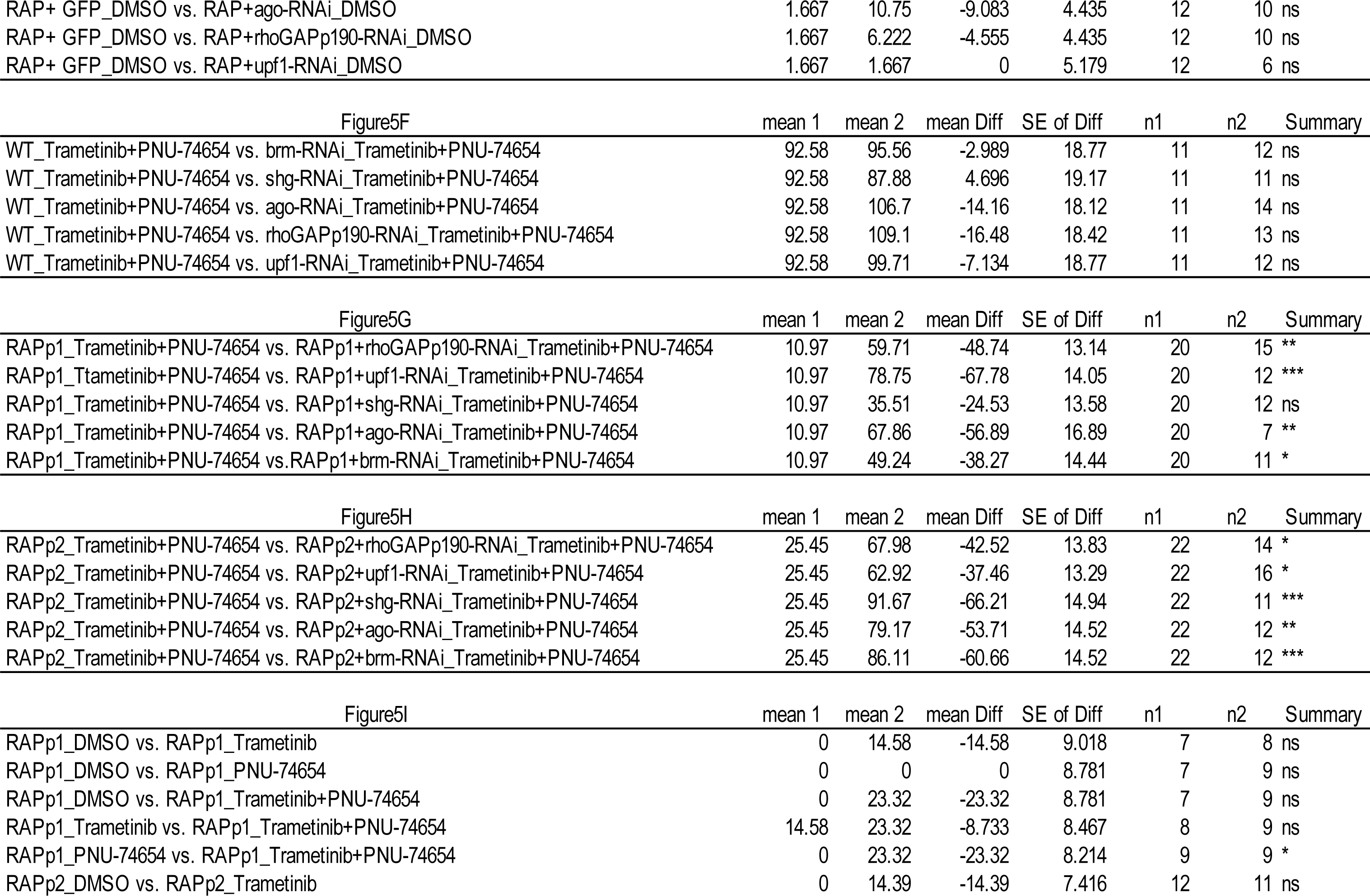
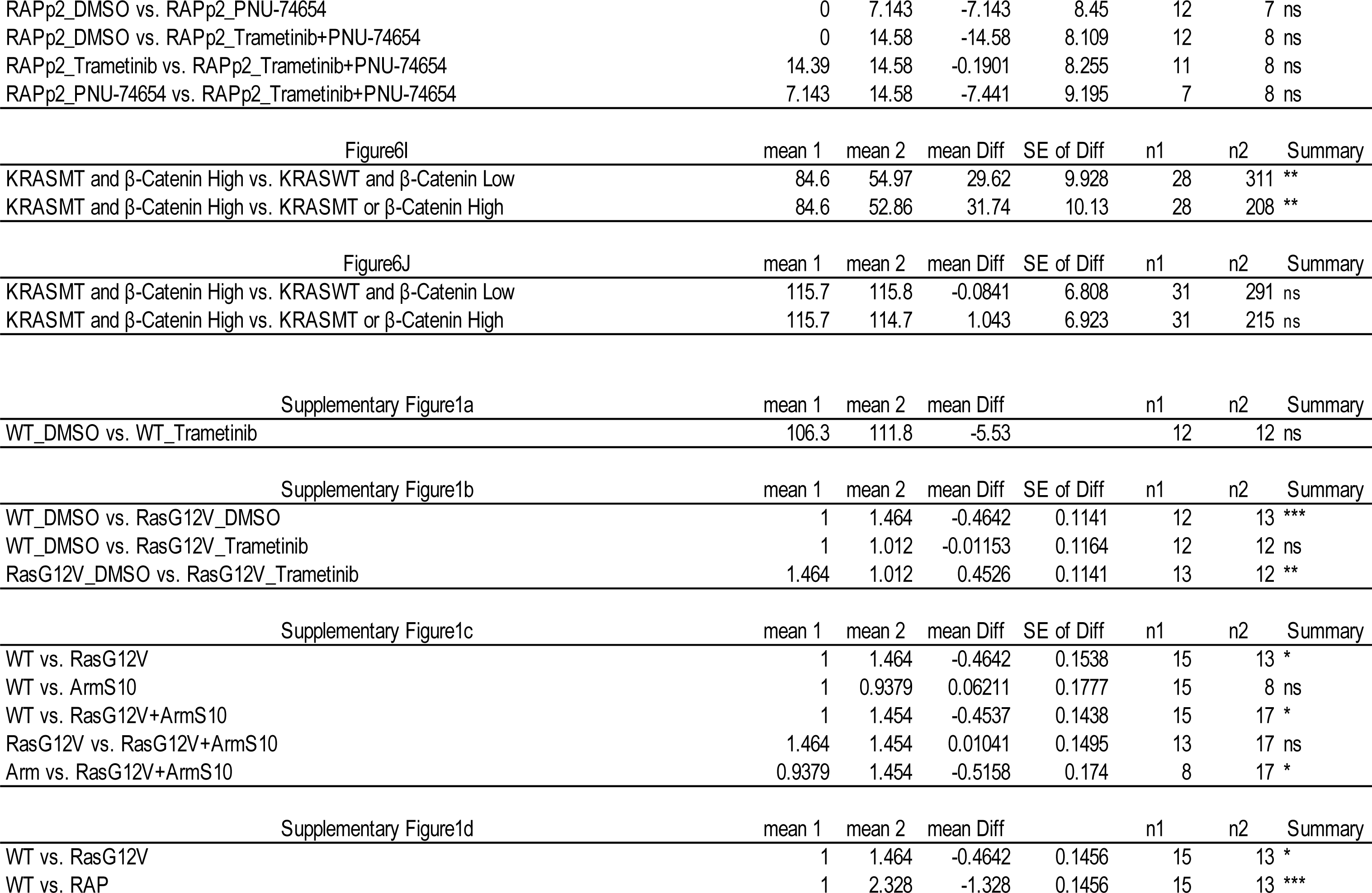
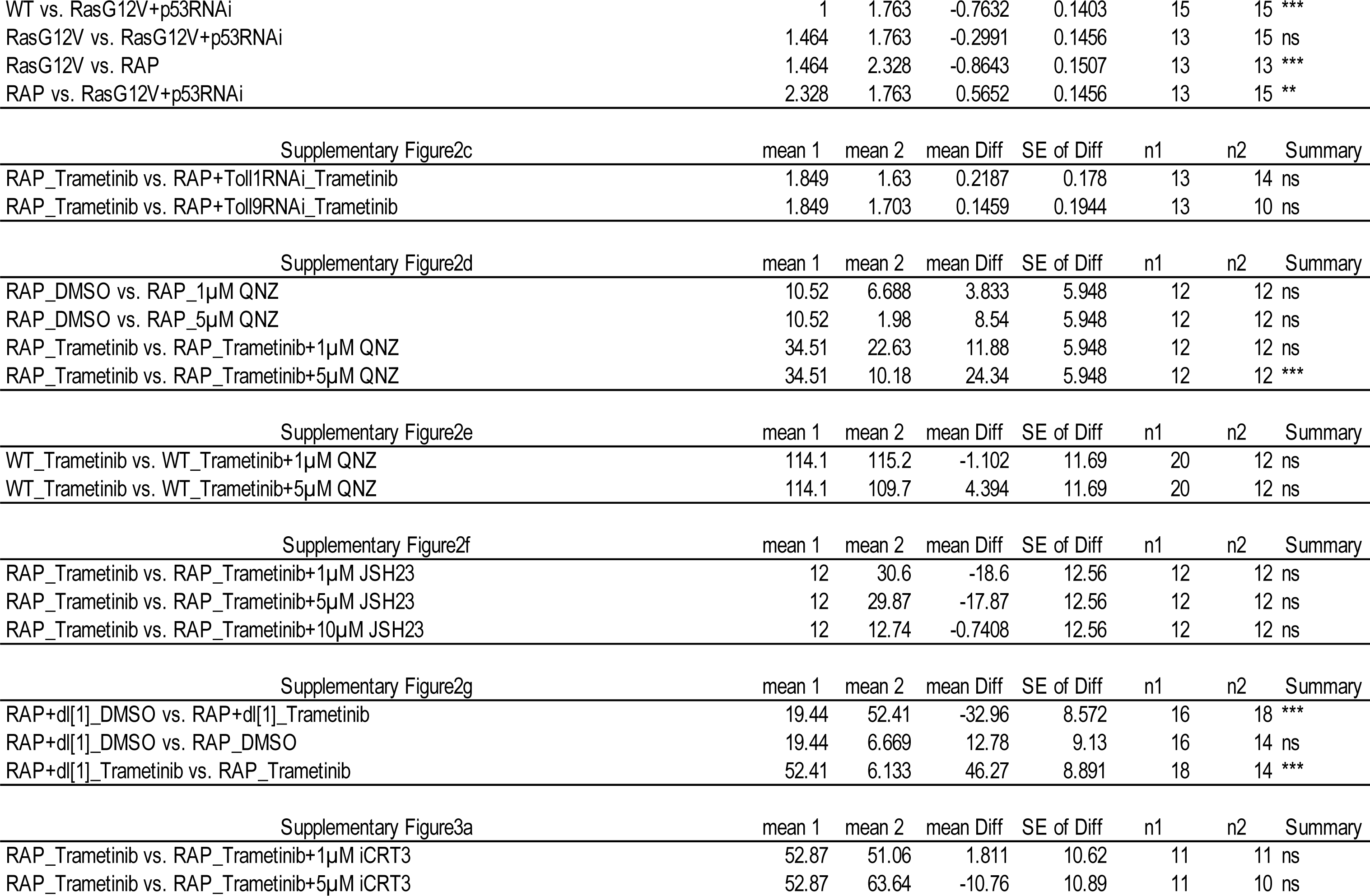
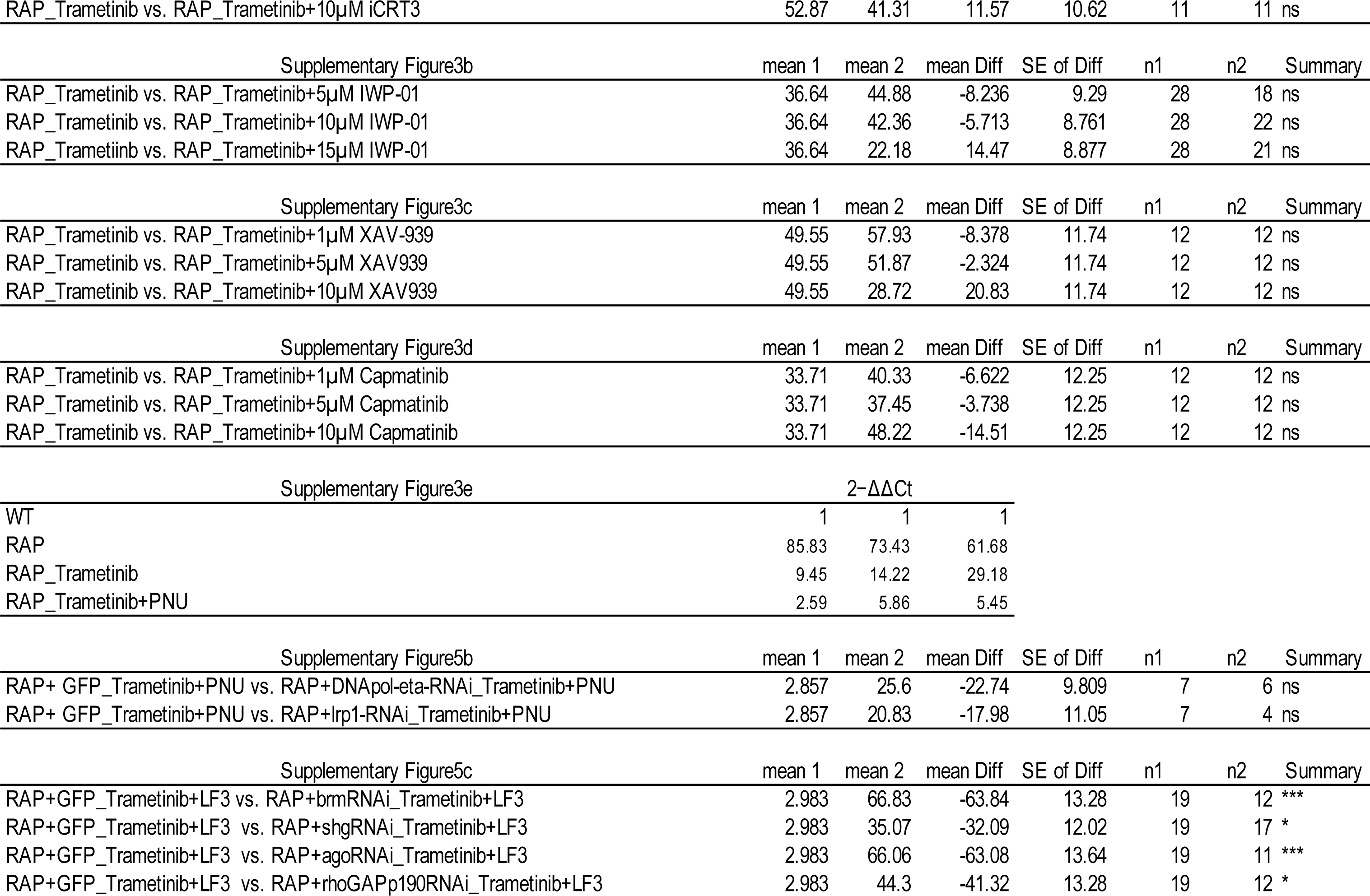
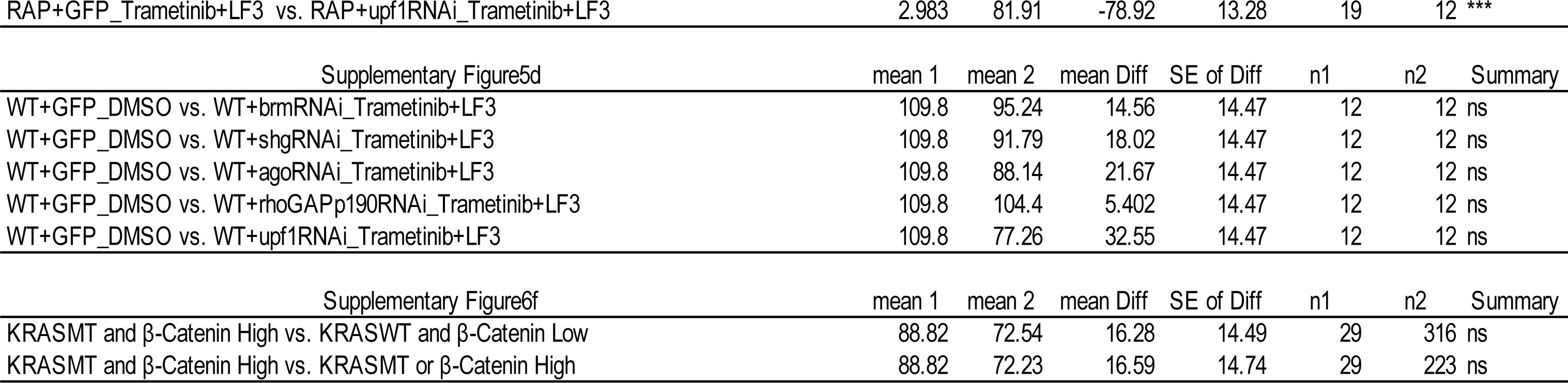

## Supplementary Table 2

### Detailed genotypes

#### Figure 1

*+/+ or Y; +/+; byn-Gal4, UAS-GFP, tub-Gal80^TS^/+* (A, B and G), *+/+ or Y; UAS-Ras^G12V^/+; byn-Gal4, UAS-GFP, tub-Gal80^TS^/+* (A and G), *+/+ or Y; UAS-Ras^G12V^, Apc-RNAi, UAS-P53-RNAi/+; byn-Gal4, UAS-GFP, tub-Gal80^TS^/+* (A and B), *+/+ or Y; UAS-Ras^G12V^, Apc-RNAi, UAS-P53-RNAi/UAS-GFP; byn-Gal4, UAS-GFP, tub-Gal80^TS^/+* (H and K), *+/+ or Y; UAS-Ras^G12V^, Apc-RNAi, UAS-P53-RNAi/+; byn-Gal4, UAS-GFP, tub-Gal80^TS^/UAS-dif-RNAi* (C, J and K), *+/+ or Y; UAS-Ras^G12V^, Apc-RNAi, UAS-P53-RNAi/UAS-dl-RNAi; byn-Gal4, UAS-GFP, tub-Gal80^TS^/+* (C, I and K), *+/+ or Y; UAS-GFP/+; byn-Gal4, UAS-GFP, tub-Gal80^TS^/+* (D and F), *+/+ or Y; +/+; byn-Gal4, UAS-GFP, tub-Gal80^TS^/UAS-dif-RNAi* (D), *+/+ or Y; UAS-dl-RNAi/+; byn-Gal4, UAS-GFP, tub-Gal80^TS^/+* (D), *+/+ or Y; UAS-Ras^G12V^/UAS-GFP; byn-Gal4, UAS-GFP, tub-Gal80^TS^/+* (E), *+/+ or Y; UAS-Ras^G12V^/UAS-cact-RNAi; byn-Gal4, UAS-GFP, tub-Gal80^TS^/+* (E), *+/+ or Y; UAS-cact-RNAi/+; byn-Gal4, UAS-GFP, tub-Gal80^TS^/+* (F), *UAS-Arm^S10^/+ or Y; +/+; byn-Gal4, UAS-GFP, tub-Gal80^TS^/+* (G), *UAS-Arm^S10^/+ or Y; UAS-Ras^G12V^/+; byn-Gal4, UAS-GFP, tub-Gal80^TS^/+* (G).

#### Figure 2

*+/+ or Y; +/+; byn-Gal4, UAS-GFP, tub-Gal80^TS^/+* (A), *+/+ or Y; UAS-Ras^G12V^, Apc-RNAi, UAS-P53-RNAi/+; byn-Gal4, UAS-GFP, tub-Gal80^TS^/+* (B), *+/+ or Y; UAS-Ras^G12V^, Apc-RNAi, UAS-P53-RNAi/+; byn-Gal4, UAS-GFP, tub-Gal80^TS^/UAS-Toll1-RNAi* (C and E), *+/+ or Y; UAS-Ras^G12V^, Apc-RNAi, UAS-P53-RNAi/+; byn-Gal4, UAS-GFP, tub-Gal80^TS^/UAS-Toll9-RNAi* (D and E), *+/+ or Y; UAS-GFP/+; byn-Gal4, UAS-GFP, tub-Gal80^TS^/+* (F), *+/+ or Y; +/+; byn-Gal4, UAS-GFP, tub-Gal80^TS^/UAS-Toll1-RNAi* (F), *+/+ or Y; +/+; byn-Gal4, UAS-GFP, tub-Gal80^TS^/UAS-Toll9-RNAi* (F).

#### Figure 3

*+/+ or Y; UAS-Ras^G12V^, Apc-RNAi, UAS-P53-RNAi/+; byn-Gal4, UAS-GFP, tub-Gal80^TS^/+* (B, C, E and F), *+/+ or Y; +/+; byn-Gal4, UAS-GFP, tub-Gal80^TS^/+* (D).

#### Figure 4

*+/+ or Y; UAS-shg-RNAi, UAS-put-RNAi, UAS-p38a-RNAi, UAS-ft-RNAi, UAS-brm-RNAi, UAS-ago-RNAi, UAS-apc-RNAi, UAS-p53-RNAi, UAS-Ras^G12V^/+; byn-Gal4, UAS-GFP, tub-Gal80^TS^/UAS-ago-RNAi, UAS-apc-RNAi* (A, CPCT006), *+/+ or Y; UAS-pten-RNAi, UAS-pten-RNAi, UAS-apc-RNAi, UAS-apc-RNAi, UAS-smox-RNAi, UAS-smox-RNAi, UAS-p53-RNAi, UAS-p53-RNAi, UAS-Ras^G12V^/+; byn-Gal4, UAS-GFP, tub-Gal80^TS^/UAS-upf1--RNAi, UAS-nej-RNAi, UASrhoGAPp190-RNAi, UAS-tefu-RNAi, UAS-apc-RNAi,UAS-smox-RNAi, UAS-pten-RNAi, UAS-p53-RNAi* (B, CPCT018), *+/+ or Y; UAS-lrp1-RNAi, UAS-dnapol-eta-RNAi, UAS-rad51c-RNAi, UAS-pc-RNAi, UAS-CG13344-RNAi, UAS-nos-RNAi, UAS-smox-RNAi, UAS-p53-RNAi, pvr/+; byn-Gal4, UAS-GFP, tub-Gal80^TS^/ UAS-Ras^G12V^, UAS-apc-RNAi, UAS-apc-RNAi, UAS-apc-RNAi, UAS-apc-RNAi, UAS-smox-RNAi, UAS-smox-RNAi, UAS-p53-RNAi, UAS-p53-RNAi* (C, CPCT045). *+/+ or Y; UAS-ird1-RNAi, UAS-wdb-RNAi, UAS-scat-RNAi, UAS-debcl-RNAi, UAS-smox-RNAi, UAS-smox-RNAi, UAS-pten-RNAi, UAS-pten-RNAi, pvr, chico/+; byn-Gal4, UAS-GFP, tub-Gal80^TS^/ UAS-Ras^G12V^, UAS-apc-RNAi, UAS-apc-RNAi, UAS-apc-RNAi, UAS-apc-RNAi, UAS-p53-RNAi, UAS-p53-RNAi, UAS-p53-RNAi, UAS-p53-RNAi* (D, CPCT050), *+/+ or Y; UAS-med-RNAi, UAS-med-RNAi, UAS-apc-RNAi, UAS-apc-RNAi, UAS-smox-RNAi, UAS-smox-RNAi, UAS-p53-RNAi, UAS-p53-RNAi, chico, UAS-Ras^G12V^/+; byn-Gal4, UAS-GFP, tub-Gal80^TS^/UAS-mus81-RNAi, UAS-pi4kIIIα-RNAi, UAS-CG4238-RNAi, UAS-fur2-RNAi, UAS-trr-RNAi, UAS-med-RNAi, UAS-p53-RNAi, UAS-apc-RNAi* (E, CPCT029).

#### Figure 5

*+/+ or Y; UAS-Ras^G12V^, Apc-RNAi, UAS-P53-RNAi/UAS-GFP; byn-Gal4, UAS-GFP, tub-Gal80^TS^/+* (A, B, D and E), *+/+ or Y; UAS-Ras^G12V^, Apc-RNAi, UAS-P53-RNAi/+; byn-Gal4, UAS-GFP, tub-Gal80^TS^/UAS-brm-RNAi* (A, B, D and E), *+/+ or Y; UAS-Ras^G12V^, Apc-RNAi, UAS-P53-RNAi/+; byn-Gal4, UAS-GFP, tub-Gal80^TS^/UAS-ago-RNAi* (A, B, D and E), *+/+ or Y; UAS-Ras^G12V^, Apc-RNAi, UAS-P53-RNAi/UAS-shg-RNAi; byn-Gal4, UAS-GFP, tub-Gal80^TS^/+* (A, B, D and E), *+/+ or Y; UAS-Ras^G12V^, Apc-RNAi, UAS-P53-RNAi/UAS-upf1-RNAi; byn-Gal4, UAS-GFP, tub-Gal80^TS^/+* (A, B, D and E), *+/+ or Y; UAS-Ras^G12V^, Apc-RNAi, UAS-P53-RNAi/UAS-rhoGAPp190-RNAi; byn-Gal4, UAS-GFP, tub-Gal80^TS^/+* (A, B, D and E), *+/+ or Y; +/UAS-GFP; byn-Gal4, UAS-GFP, tub-Gal80^TS^/+* (F), *+/+ or Y; +/+; byn-Gal4, UAS-GFP, tub-Gal80^TS^/UAS-brm-RNAi* (F), *+/+ or Y; +/+; byn-Gal4, UAS-GFP, tub-Gal80^TS^/UAS-ago-RNAi* (F), *+/+ or Y; +/UAS-shg-RNAi; byn-Gal4, UAS-GFP, tub-Gal80^TS^/+* (F), *+/+ or Y; +/UAS-upf1-RNAi; byn-Gal4, UAS-GFP, tub-Gal80^TS^/+* (F), *+/+ or Y; +/UAS-rhoGAPp190-RNAi; byn-Gal4, UAS-GFP, tub-Gal80^TS^/+* (F), *+/+ or Y; UAS-ago-RNAi, UAS-wts-RNAi, UAS-CG7742-RNAi, UAS-Atg2-RNAi, UAS-Ras^G12V^, Apc-RNAi, UAS-P53-RNAi/UAS-GFP; byn-Gal4, UAS-GFP, tub-Gal80^TS^/+* (RAPp1, G and I), *+/+ or Y; UAS-ago-RNAi, UAS-wts-RNAi, UAS-CG7742-RNAi, UAS-Atg2-RNAi, UAS-Ras^G12V^, Apc-RNAi, UAS-P53-RNAi/+; byn-Gal4, UAS-GFP, tub-Gal80^TS^/UAS-brm-RNAi* (RAPp1, G and I), *+/+ or Y; UAS-ago-RNAi, UAS-wts-RNAi, UAS-CG7742-RNAi, UAS-Atg2-RNAi, UAS-Ras^G12V^, Apc-RNAi, UAS-P53-RNAi/+; byn-Gal4, UAS-GFP, tub-Gal80^TS^/UAS-ago-RNAi* (RAPp1, G and I), *+/+ or Y; UAS-ago-RNAi, UAS-wts-RNAi, UAS-CG7742-RNAi, UAS-Atg2-RNAi, UAS-Ras^G12V^, Apc-RNAi, UAS-P53-RNAi/UAS-shg-RNAi; byn-Gal4, UAS-GFP, tub-Gal80^TS^/+* (RAPp1, G and I), *+/+ or Y; UAS-ago-RNAi, UAS-wts-RNAi, UAS-CG7742-RNAi, UAS-Atg2-RNAi, UAS-Ras^G12V^, Apc-RNAi, UAS-P53-RNAi/UAS-upf1-RNAi; byn-Gal4, UAS-GFP, tub-Gal80^TS^/+* (RAPp1, G and I), *+/+ or Y; UAS-ago-RNAi, UAS-wts-RNAi, UAS-CG7742-RNAi, UAS-Atg2-RNAi, UAS-Ras^G12V^, Apc-RNAi, UAS-P53-RNAi/UAS-rhoGAPp190-RNAi; byn-Gal4, UAS-GFP, tub-Gal80^TS^/+* (RAPp1, G and I), *+/+ or Y; UAS-vrp1-RNAi, UAS-ry-RNAi, UAS-khc-73-RNAi, UAS-Ras^G12V^, Apc-RNAi, UAS-P53-RNAi/UAS-GFP; byn-Gal4, UAS-GFP, tub-Gal80^TS^/+* (RAPp2, H and I), *+/+ or Y; UAS-vrp1-RNAi, UAS-ry-RNAi, UAS-khc-73-RNAi, UAS-Ras^G12V^, Apc-RNAi, UAS-P53-RNAi/+; byn-Gal4, UAS-GFP, tub-Gal80^TS^/UAS-brm-RNAi* (RAPp2, H and I), *+/+ or Y; UAS-vrp1-RNAi, UAS-ry-RNAi, UAS-khc-73-RNAi, UAS-Ras^G12V^, Apc-RNAi, UAS-P53-RNAi/+; byn-Gal4, UAS-GFP, tub-Gal80^TS^/UAS-ago-RNAi* (RAPp2, H and I), *+/+ or Y; UAS-vrp1-RNAi, UAS-ry-RNAi, UAS-khc-73-RNAi, UAS-Ras^G12V^, Apc-RNAi, UAS-P53-RNAi/UAS-shg-RNAi; byn-Gal4, UAS-GFP, tub-Gal80^TS^/+* (RAPp2, H and I), *+/+ or Y; UAS-vrp1-RNAi, UAS-ry-RNAi, UAS-khc-73-RNAi, UAS-Ras^G12V^, Apc-RNAi, UAS-P53-RNAi/UAS-upf1-RNAi; byn-Gal4, UAS-GFP, tub-Gal80^TS^/+* (RAPp2, H and I), *+/+ or Y; UAS-vrp1-RNAi, UAS-ry-RNAi, UAS-khc-73-RNAi, UAS-Ras^G12V^, Apc-RNAi, UAS-P53-RNAi/UAS-rhoGAPp190-RNAi; byn-Gal4, UAS-GFP, tub-Gal80^TS^/+* (RAPp2, H and I).

#### Supplementary Figure 1

*+/+ or Y; +/+; byn-Gal4, UAS-GFP, tub-Gal80^TS^/+* (a-e), *+/+ or Y; UAS-Ras^G12V^/+; byn-Gal4, UAS-GFP, tub-Gal80^TS^/+* (b, c, d, f and g), *UAS-Arm^S10^/+ or Y; +/+; byn-Gal4, UAS-GFP, tub-Gal80^TS^/+* (c and h), *UAS-Arm^S10^/+ or Y; UAS-Ras^G12V^/+; byn-Gal4, UAS-GFP, tub-Gal80^TS^/+* (c and i), *+/+ or Y; UAS-Ras^G12V^, Apc-RNAi, UAS-P53-RNAi/+; byn-Gal4, UAS-GFP, tub-Gal80^TS^/+* (d and k). *+/+ or Y; UAS-Ras^G12V^, UAS-P53-RNAi/+; byn-Gal4, UAS-GFP, tub-Gal80^TS^/+* (d and j).

#### Supplementary Figure 2

*+/+ or Y; UAS-Ras^G12V^, Apc-RNAi, UAS-P53-RNAi/UAS-GFP; byn-Gal4, UAS-GFP, tub-Gal80^TS^/+* (c), *+/+ or Y; UAS-Ras^G12V^, Apc-RNAi, UAS-P53-RNAi/+; byn-Gal4, UAS-GFP, tub-Gal80^TS^/UAS-Toll1-RNAi* (a and c), *+/+ or Y; UAS-Ras^G12V^, Apc-RNAi, UAS-P53-RNAi/+; byn-Gal4, UAS-GFP, tub-Gal80^TS^/UAS-Toll9-RNAi* (b and c), *+/+ or Y; UAS-Ras^G12V^, Apc-RNAi, UAS-P53-RNAi/+; byn-Gal4, UAS-GFP, tub-Gal80^TS^/+* (d, f and g), *+/+ or Y; +/+; byn-Gal4, UAS-GFP, tub-Gal80^TS^/+* (e), *+/+ or Y; UAS-Ras^G12V^, Apc-RNAi, UAS-P53-RNAi/dl[1]; byn-Gal4, UAS-GFP, tub-Gal80^TS^/+* (g).

#### Supplementary Figure 3

*+/+ or Y; UAS-Ras^G12V^, Apc-RNAi, UAS-P53-RNAi/+; byn-Gal4, UAS-GFP, tub-Gal80^TS^/+* (a-e), *+/+ or Y; +/+; byn-Gal4, UAS-GFP, tub-Gal80^TS^/+* (e).

#### Supplementary Figure 5

*+/+ or Y; UAS-Ras^G12V^, Apc-RNAi, UAS-P53-RNAi/UAS-GFP; byn-Gal4, UAS-GFP, tub-Gal80^TS^/+* (b and c), *+/+ or Y; UAS-Ras^G12V^, Apc-RNAi, UAS-P53-RNAi/+; byn-Gal4, UAS-GFP, tub-Gal80^TS^/UAS-DNApol-eta-RNAi* (b and c), *+/+ or Y; UAS-Ras^G12V^, Apc-RNAi, UAS-P53-RNAi/+; byn-Gal4, UAS-GFP, tub-Gal80^TS^/UAS-lrp1-RNAi* (b and c), *+/+ or Y; UAS-Ras^G12V^, Apc-RNAi, UAS-P53-RNAi/UAS-tefu-RNAi; byn-Gal4, UAS-GFP, tub-Gal80^TS^/+* (b and c), *+/+ or Y; UAS-Ras^G12V^, Apc-RNAi, UAS-P53-RNAi/+; byn-Gal4, UAS-GFP, tub-Gal80^TS^/UAS-nej-RNAi* (b and c), *+/+ or Y; UAS-Ras^G12V^, Apc-RNAi, UAS-P53-RNAi/+; byn-Gal4, UAS-GFP, tub-Gal80^TS^/UAS-nos-RNAi* (b and c), *+/+ or Y; UAS-Ras^G12V^, Apc-RNAi, UAS-P53-RNAi/UAS-pc-RNAi; byn-Gal4, UAS-GFP, tub-Gal80^TS^/+* (b and c), *+/+ or Y; UAS-Ras^G12V^, Apc-RNAi, UAS-P53-RNAi/UAS-rad51C-RNAi; byn-Gal4, UAS-GFP, tub-Gal80^TS^/+* (b and c), *+/+ or Y; +/UAS-GFP; byn-Gal4, UAS-GFP, tub-Gal80^TS^/+* (d), *+/+ or Y; +/+; byn-Gal4, UAS-GFP, tub-Gal80^TS^/UAS-brm-RNAi* (d), *+/+ or Y; +/+; byn-Gal4, UAS-GFP, tub-Gal80^TS^/UAS-ago-RNAi* (d), *+/+ or Y; +/UAS-shg-RNAi; byn-Gal4, UAS-GFP, tub-Gal80^TS^/+* (d), *+/+ or Y; +/UAS-upf1-RNAi; byn-Gal4, UAS-GFP, tub-Gal80^TS^/+* (d), *+/+ or Y; +/UAS-rhoGAPp190-RNAi; byn-Gal4, UAS-GFP, tub-Gal80^TS^/+* (d).

